# PHLOWER - Single cell trajectory analysis using Hodge Decomposition

**DOI:** 10.1101/2024.10.01.613179

**Authors:** Mingbo Cheng, Jitske Jansen, Katharina Reimer, Vincent Grande, James Shiniti Nagai, Zhijian Li, Paul Kießling, Martin Grasshoff, Christoph Kuppe, Michael T. Schaub, Rafael Kramann, Ivan G. Costa

## Abstract

Multi-modal single-cell sequencing, which captures changes in chromatin and gene expression in the same cells, is a game changer in the study of gene regulation in cellular differentiation processes. Computational trajectory analysis is a key computational task for inferring differentiation trees from this single-cell data, though current methods struggle with complex, multi-branching trees and multi-modal data. To address this, PHLOWER (decomPosition of the Hodge Laplacian for inferring trajectOries from floWs of cEll diffeRentiation) leverages the harmonic component of the Hodge decomposition on simplicial complexes to infer trajectory embeddings. These natural representations of cell differentiation facilitate the estimation of their underlying differentiation trees. We evaluate PHLOWER through benchmarking with multi-branching differentiation trees and using novel kidney organoid multi-modal and spatial single-cell data. These demonstrate the power of PHLOWER in both the inference of complex trees and the identification of transcription factors regulating off-target cells in kidney organoids.

## 1 Introduction

Cellular differentiation, the process by which a cell changes its chromatin and expression programs to acquire more specialized functions, is not only crucial in the development of multicellular organisms but also key during onset and progression of diseases. Transcription factors, which are proteins binding to regulatory DNA regions (open chromatin regions), are key regulators of gene expression thereby orchestrating cellular differentiation processes. Dissecting these key regulatory events will help to develop protocols for cellular reprogramming or ex vivo differentiation in e.g. organoids and to understand disease-related differentiation processes for potential therapeutic interventions^1^. In this context, a unique resource to understand the interplay between chromatin, regulatory signals (transcription factor binding), and expression changes during cellular differentiation^2,3^, is multi-modal single-cell sequencing^2^, which can measure both full expression programs and genome-wide open chromatin. However, the associated experimental protocols dissociate cells, making it impossible to track how an individual cell differentiates over time experimentally.

To overcome this experimental limitation, computational trajectory analysis, which explores non-linear embeddings in the cellular space and algorithms to find trees in these spaces, has therefore become an important tool in the analysis of cell differentiation in single cells^4–7^. However, these approaches have been hitherto applied to the study of differentiation trees with small trees (3-9 branches) and the only comprehensive benchmarking is based on small trees with four to five differentiation branches^8^. Currently, no work has evaluated their scalability to complex and multi-branching trajectories. Moreover, the analysis of single-cell multiomics data provides further challenges, as there is a need to estimate joint embeddings across modalities for modality detection^9,10^. Altogether, there is a lack of computational approaches to infer complex, branching trajectories from multimodal sequencing data.

We propose here PHLOWER (*decom***P**osition of the **H**odge **L**aplacian for inferring traject**O**ries from flo**W**s of c**E**ll diffe**R**entiation). Simplicial complexes (SCs) are generalizations of graphs that allow not only for nodes and edges, but also triangles and other higher-order structures to be present^11^. The discrete Hodge Laplacian (HL) on simplical complexes represent a generalization of the well-known graph Laplacian explored in diffusion maps^12^ and trajectory inference for single cell data^13,14^. PHLOWER uses SCs as representation of single cell multimodal data, where a node represents a cell and an edge indicates a potential cell differentiation event. The harmonic component obtained via spectral decomposition of the Hodge Laplacian on a SC allow the creation of edge-flow embeddings (cell differentiation events) and trajectory embeddings (cell differentiation paths)^11^. These represent cell differentiation processes directly, and can thus enable the detection of complex differentiation trees and characterize branching events with high precision.

## 2 Results

### 2.1 Differentiation Tree Inference with PHLOWER

PHLOWER uses the discrete Hodge Laplacian (HL) and its associated Hodge decomposition to obtain embeddings of cell differentiation trajectories. The Graph Laplacian (a zero-order Hodge Laplacian) is a matrix representation of graphs, where samples are encoded as vertices and distances as edge weights. Particular eigenvectors of the Graph Laplacian are used to obtain an embedding representing cells in the graph, forming the basis for methods in clustering analysis^17^, non-linear dimension reduction^12^, trajectory inference and pseudo-time estimation in single-cell data^13,14^. In PHLOWER, we represent single cell data as a simplicial complex, i.e., a higher-order generalization of a graph, consisting of nodes (0-simplices), edges (1-simplices) and triangles (2-simplices). The spectral decomposition of the HL can be used to decompose edge-flows on into gradient-free, curl-free, and harmonic components^11,18^. Of note, while the Hodge Laplacian and Hodge decomposition are defined on differential forms on Riemannian manifolds^19^, there are guarantees that the behavior of the Hodge Laplacian on a simplicial complex converges to the Hodge Laplacian on manifolds in the limit^20,21^. PHLOWER focuses on the harmonic eigenvectors of the HL^11,18^ as these are associated to holes in the simplicial complex, which in turn can reveal cell differentiation tree branches in the gene expression SC.

In short, PHLOWER first uses a graph representation of the single cell data and the graph Laplacian to estimate pseudo-time of cells and identify progenitor and terminal cells similar as before^13,22^. Cell differentiation processes are classically represented as generating an out-branching, tree-like structure. To transform such a directed branching process into a simplicial complex, we perform a Delaunay triangulation and connect terminal differentiated cells (high pseudo-time) with progenitor cells (low pseudo-time). Connecting cells with high and low pseudotime creates a hole for every main trajectory in the graph (Fig. 1). Next, we perform a Hodge Decomposition of the edge flow

**Figure 1.**
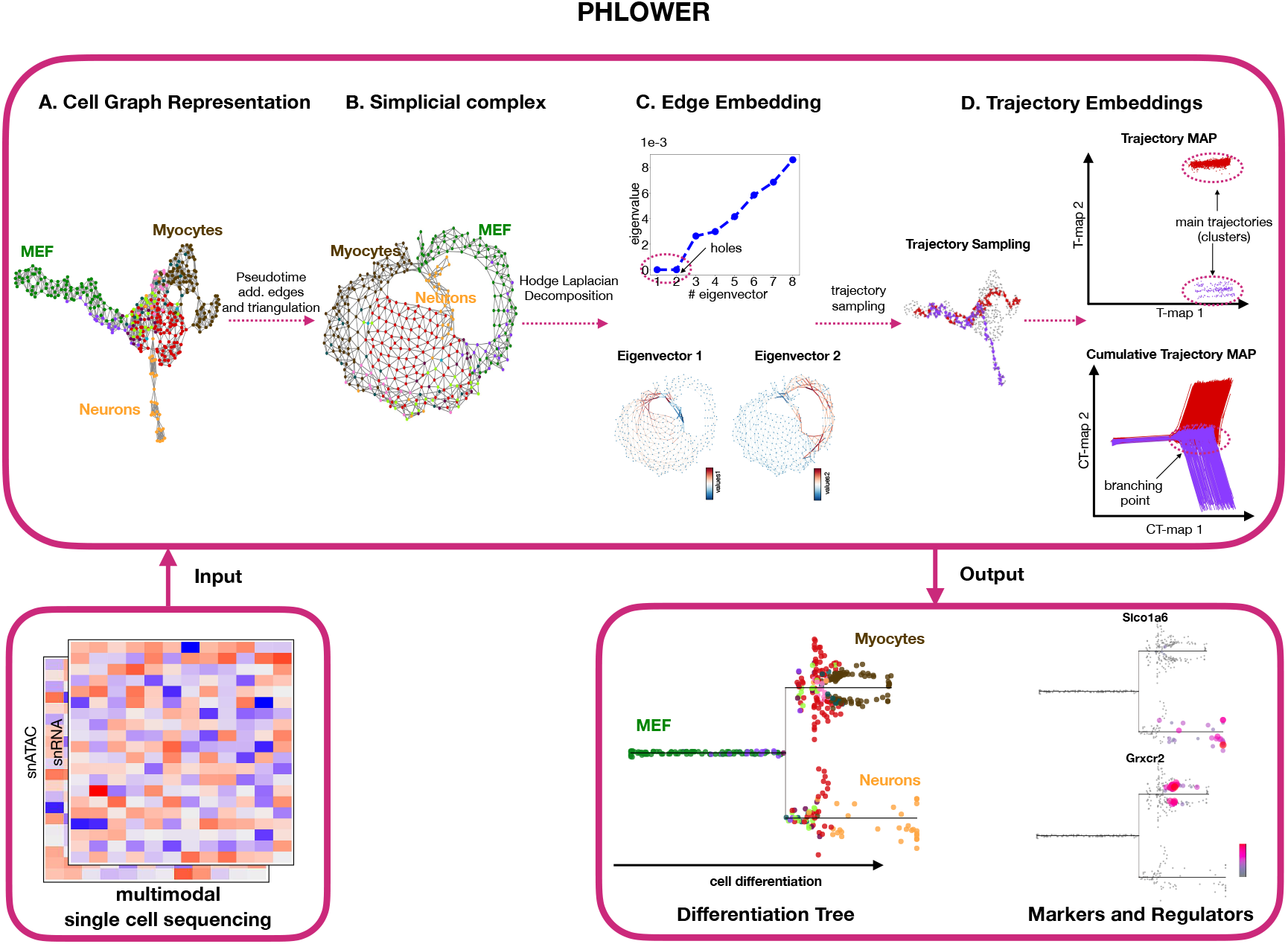
Schematic overview of major steps of PHLOWER. PHLOWER receives as input a multimodal or an unimodal single cell dataset. **(A)** It next creates a graph representation and a graph embedding^15^ of the cells. Our example is based on a differentiation system of mouse embryonic fibroblasts (MEF) towards neurons or myocytes^16^. From this, PHLOWER uses a Graph Laplacian decomposition and random-walk to estimate pseudo-time from progenitor cells (i.e. MEF). **(B)**, Next, cells with low pseudotime (progenitors) and high pseudotime (terminally differentiated cells) are connected and a simplicial complex (SC) is obtained by Delaunay triangulation of the graph. **(C)** An edge embedding is obtained via the harmonic eigenvectors via the decomposition of the SC Hodge Laplacian. In the MEF data, the first two eigenvectors have zero eigenvalues and their signals discriminate edges belonging to the two differentiation trajectories (or holes in the SC). **(D)** PHLOWER performs next random walks to obtain trajectories and uses the HL decomposition to obtain a trajectory embedding and a cumulative trajectory embedding. Clustering analysis in the trajectory embedding reveals major trajectories in the data, while the cumulative embedding space is used to estimate trajectory backbones and branching point events. PHLOWER outputs stream trees representing the differentiation tree over pseudo-time and allow the detection of regulators and gene markers.

space on the SC. The harmonic components of this decomposition provide edge level embeddings, where each point in an embedding represents a cell differentiation event (Eq. 19), as well as a trajectory embedding, where each embedding point represents a cell differentiation trajectory (Eq. 21). PHLOWER explores these embeddings to delineate major differentiation trajectories in single-cell data and to reconstruct complex cell differentiation trees. As an example, we show how PHLOWER can be used to predict a simple differentiation tree from mouse embryonic cells (MEF)^16^ towards either neurons or myocytes (Fig. 1 and Supp. Fig. S1).

### Benchmarking of Trajectory Inference Methods

Next, we evaluated PHLOWER and competing methods on simulated datasets^23^ and scRNA-seq data-sets^24^ on how well methods can recover original tree structures and how well they can place cells within these differentiation trees. We utilized the diffusion-limited aggregation tree (DLA tree)^23^ to simulate ten complex differentiation tree dataset with 5 to 18 total branches similar as in^14^. We selected 20 scRNA-seq datasets from the benchmarking Dynverse, which contained tree structures with at least three branches (Supp. Table S1).

These data were provided as input for PHLOWER and competing approaches for the detection of differentiation trees (PAGA tree^4^, monocle 3^25^, cellTree^26^, pCreode^27^, Slice^28^, RaceID/StemID^29,30^, slingshot^6^, TSCAN^31^, Slicer^32^, MST^33^, Elpigraph^34^, and STREAM^5^). This selection included the top 10 approaches from a recent benchmark evaluation^24^ ^1^. Tools were evaluated using the metrics proposed by the Dynverse framework^24^: tree structure similarity (Hamming-Ipsen-Mikhailov; HIM), location of cells within a branch (Correlation), cell allocation to branches (F1 branches), and cell allocation to branching points (F1 milestones). An example of the steps of the PHLOWER algorithm in fitting a differentiation tree in a simulated data is shown in Supp. Fig. S2.

For simulated data, PHLOWER was the best-performing method in regard to tree topology recovery, followed by PAGA, RaceID/StemID and monocle3 (Fig.2A and Supp. Fig. S3). In the problem of allocating cells to positions in a branch, PHLOWER was also the best performer, followed by TSCAN and RaceID (Fig.2B). In the problem of allocating cells to the correct branches, PHLOWER obtained the best performance, followed by monocle3 and PAGA. Finally, in the allocation of cells to branches, PHLOWER was the best performer, followed by RaceID/StemID, monocle3, and PAGA. For scRNA-seq data, PHLOWER was the best performing approach followed by monocle3, stream and PAGA regarding structure similarity (Fig. 2E and Supp. Fig. S3). Regarding the location of cells (Fig. 2F), PHLOWER obtained highest scores followed by pcreode, slice and slingshot. Finally, regarding allocation to branches or millstones (Fig. 2G-H), PHLOWER was also the top performer with slingshot, monocle3 and slice as follow up approaches.. Altogether, PHLOWER was ranked as the top performer in all evaluated scenarios. Interestingly, follow up methods had distinct performances on simulated or scRNA-seq datasets, while raceid, monocle3 and paga were consistently well on simulated data; there was no clear consistent method scoring well on all problems in the scRNA-seq data.

**Figure 2.**
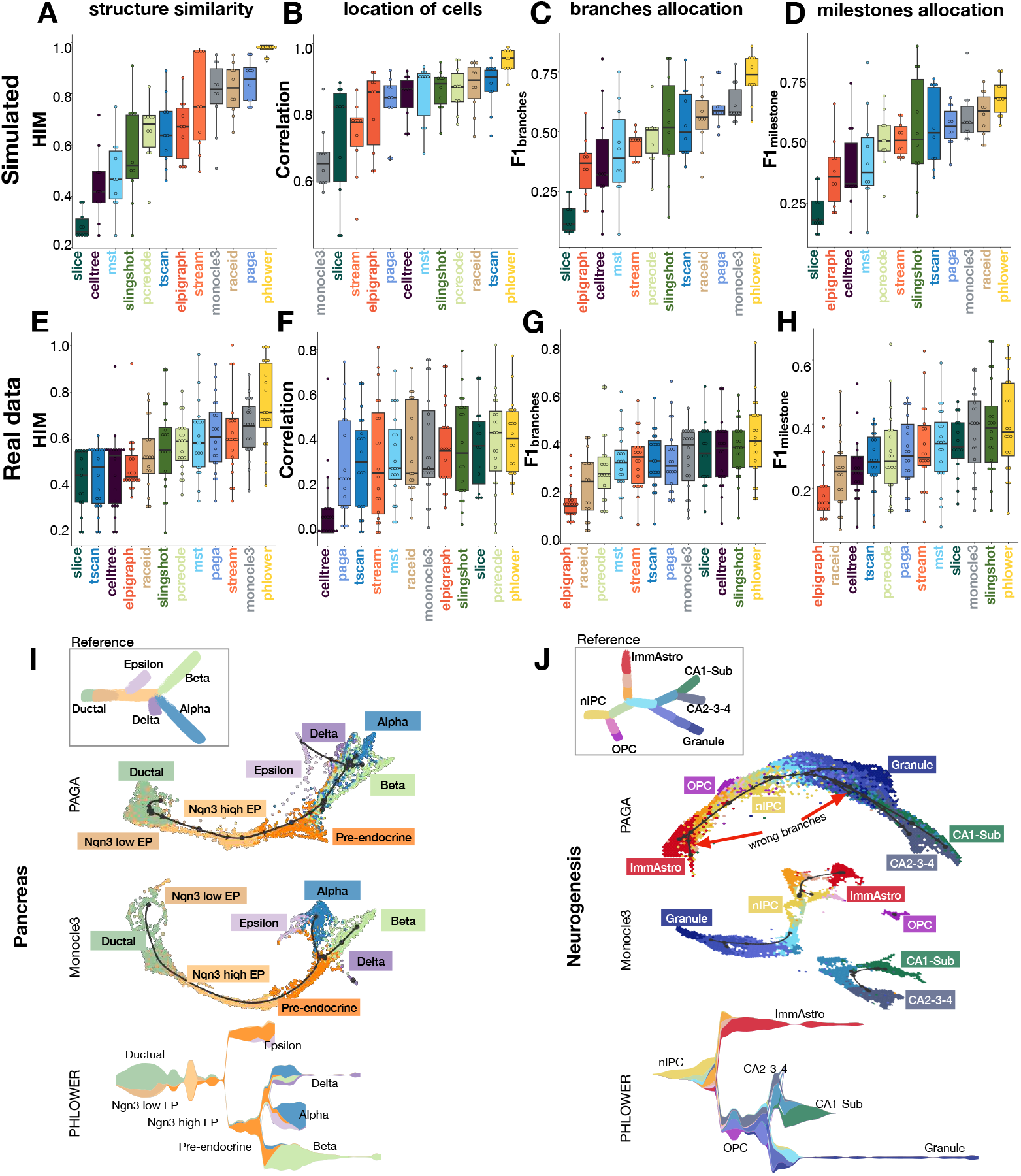
PHLOWER benchmarking. **A)** We show the HIM score (y-axis) for all 12 tree inference algorithms (x-axis) measuring topology similarity between true and inferred trees. Methods are ranked (left to right) regarding the mean value. **B-D)** Same as **A** cell allocation within a branch (correlation); Cell allocation to branching points (F1 milestones) and cell allocation to branches (F1 branches). **E-H)** We show the HIM, Correlation, F1 branches and F1 milestone scores (y-axis) for 20 dynverse tree inference algorithms (x-axis). **I-J)** Differentiation trees estimated with PAGA, Monocle3 and PHLOWER algorithms for the pancreatic progenitor and neurogenesis data sets. Trees in the top left display the reference cell differentiation tree.

Another important aspect is the computational requirements of the approaches. To evaluate this, we resorted to two data sets commonly used in the RNA velocity literature: the pancreas progenitor (≈ 3.700 cells)^36^ and neurogenesis (≈ 18.000 cells)^37^. PHLOWER required between 0.5-12 hours and 12GB-40GB of memory for these data sets using an standard desktop (Supp. Fig. S4A-B). We next evaluate an time-efficient variant of PHLOWER based on cell down-sampling (Supp. Fig. S5). We observe that by using 30% of the cells, PHLOWER obtains a 8.6x speed up and one sixth of memory in contrast to using all cells, while infering a bona-fide neurogenesis cell differentiation tree.

The performance of PHLOWER and competing methods is further illustrated by the analysis of the inferred trees in the pancreas progenitor and neurogenesis data sets. We focus here on PHLOWER, PAGE and monocle3. These were among the best performing in structure similarity benchmarking (Fig. 2A-E), and recovered more complex trees on these data sets, when compared to other approaches (Supp. Figs. S6 and S7). For pancreas progenitor data (Fig. 2I), we observed that monocle3 tree does not capture the epsilon branch, and the delta branch corresponds to an unconnected single branched tree. PHLOWER and PAGA recovered all main branches of this tree: epsilon, delta, alpha and beta. In the larger and more complex neurogenesis data Fig. 2J, monocle3 inferred 4 unconnected trees delineating some of the main branching events. PHLOWER and PAGA recapitulated the main terminal branches with the exception of the small OPC population, which was missed by both methods. Note however that PAGA inferred two false positive branches (see red arrows Fig. 2J), which are not related to any cell type described in this data set^38^. Altogether, this analysis supports the power of PHLOWER in recovery on complex cell differentiation trees.

### Inferring cellular trajectories in kidney organoids

Induced pluripotent stem cell (iPSC) derived kidney organoids represent a solid model to validate cellular trajectories since this model represents the differentiation of stem and progenitor cells towards various kidney cell lineages. We therefore generated a multimodal single cell multiome sequencing (RNA and ATAC) data sets of kidney organoids after 7, 12, 19 and 25 days of differentiation by using an own protocol^39^. This recovered 13,751 cells with an average of 10,378 RNA transcripts and 19,263 DNA fragments (ATAC) per cell (Supp. Fig. S8). We next integrated the data for each modality independently^40^ and used MOJITOO to obtain a joint ATAC-RNA embedding^9^. We provided the data as input for PHLOWER, which recovered a trajectory with nine branches (Fig. 3A; Supp. Fig. S9). We observed that the tree successfully sorted cells based on the organoids age (Fig. 3B) and this matched well with the pseudo-time estimates (Fig. 3C). These nine branches could be grouped into three major branches associated with epithelial cells (two sub-branches associated with podocytes and one with tubular cells); four branches of stromal cells and one major branch associated with muscle and neuronal cells (3 sub-branches). The annotation of these cells was based on the expression of markers as TBXT^41^ and KDR^42^ for mesoderm cells; PODXL^43^ and NPHS2^43^ for podocytes, SLC12A1^44^ and PAPPA2^44^ for kidney tubule epithelial; COL1A2^43^ and PDGFRA^45^ for stromal cells; and neuronal and muscle markers MAP2^39^, MSX1^41,43–45^, MYL1 and MYF6 (Supp. Fig. S10). The latter branches, i.e. neuronal and muscle, are considered off-target cells and potentially may hamper maturation of kidney cells in the organoids, which should be avoided in kidney organoids^46,47^.

**Figure 3.**
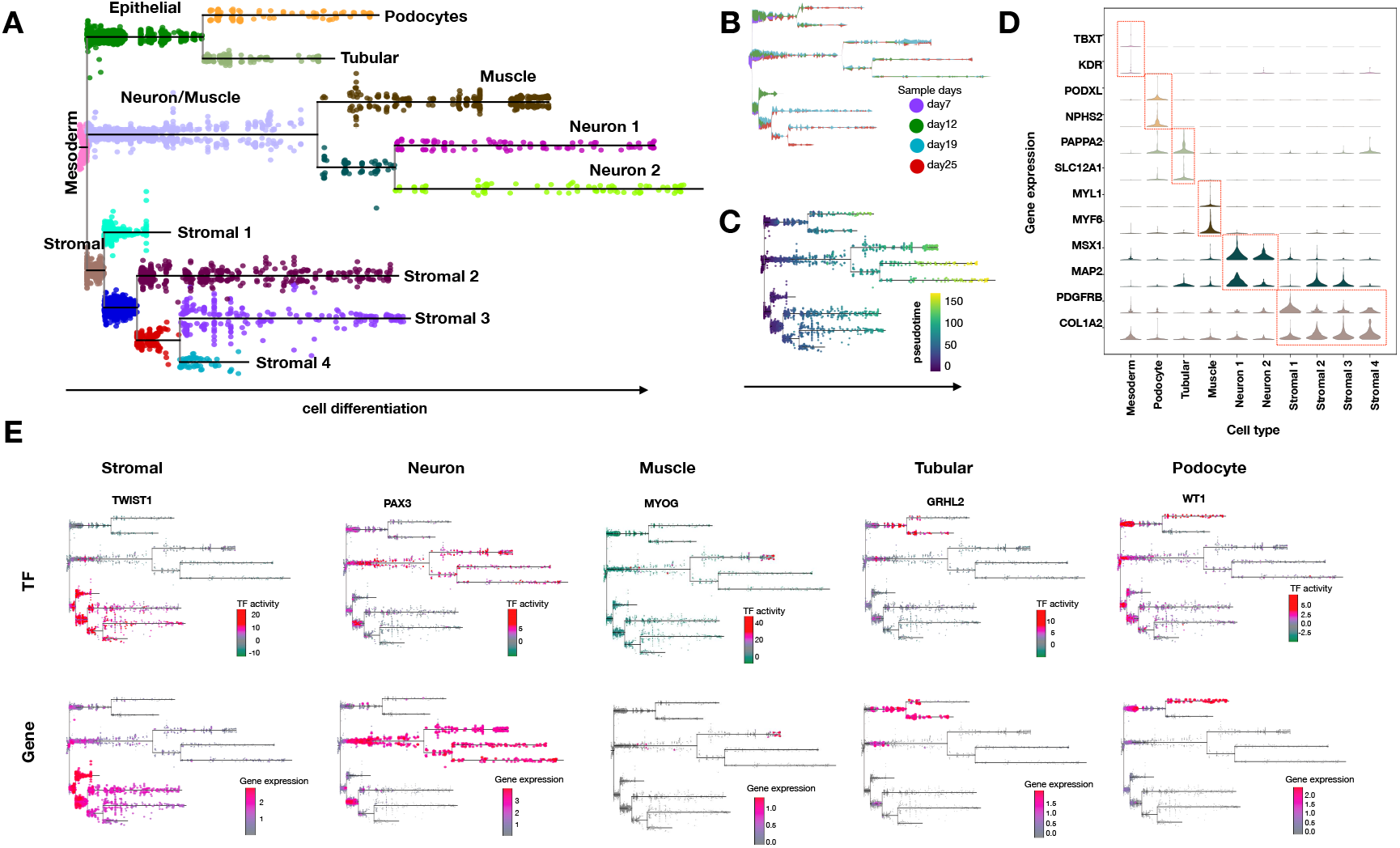
Kidney organoid data. **A)** Differentiation tree on the kidney organoid data as estimated by PHLOWER. **B-C)** Day of differentiation and pseudo-time estimates of cells in the differentiation trees. **D)** Violin plots show the marker genes for cell types in each branch. **E)** We show selected TFs with branch specific expression and TF activity scores.

A key question to be addressed by PHLOWER is the detection of regulators (transcription factors; TF) driving the differentiation of iPSC within the kidney organoids. We leveraged the tree inferred by PHLOWER and a procedure similar to scMega^48^ to detect TFs related to branch differentiation. In short, we estimated TF activity scores with ChromVAR^49^ and selected TFs, which expression are concordant with the TF activity and are differentially expressed between compared branches (Supp. Fig. S11). When comparing cells at the end of tubular and podocyte trajectories, this recovered bonafide regulators of these cells such as WT1^50^ and MAFB^51^ for podocytes and HNF1B^52^ and GRHL2^53^ for tubular cells (Fig. 3E and Supp. Fig. S12). Next, we compared major branches: stromal cells vs. others and neuronal/muscle cells vs. others. We detected TFs related to fibroblasts TWIST1 and RUNX2^54,55^ as regulators of stromal cells, and the known skeletal muscle MYOG for the muscle branch. Among the top three neuronal cells regulators, we observed the TFs PAX3, RFX4 and ZIC2, which have been previously related to neuronal cell differentiation and/or neuronal diseases^56–58^. The modulation of these TFs is of particular interest, as neuronal cells are considered to be undesired off-target cell types in kidney organoids.

### Spatial organization of kidney organoids and perturbation of off-target lineages

We leveraged our analysis of multimodal single cell data to perform a subcellular resolution spatial transcriptomic analysis of kidney organoids using the Xenium spatial imaging platform^59^. We used PHLOWER results to derive a 100 gene kidney organoid marker panel for the detection of mesenchymal, tubule-epithelial, stromal, neuronal cells and podocytes (Sup. Fig. S13A). This panel selection included computational data driven based marker genes from the multiome analysis and some literature based marker genes; and top candidate TFs from the scMega analysis (Supp. Table S2).

Clustering and trajectory analyses of the 105,092 cells in a 19 day kidney organoid and two 25 days kidney organoids indicated the identification of progenitor cells (mesoderm progenitors podocyte and tubule-epithelial progenitors) and all major kidney organoid branches (mesoderm, podocytes, tubule-epithelial-cells, stromal and neuronal-cells) (Fig. 4A-B; Supp. Fig. S13C). When contrasting cells detected on 19 vs. 25 days, we observed a higher proportion of mesoderm cells in day 19; while day 25 had a higher amount of tubule-epithelial cells and podocytes (Fig. 4C)reflecting the higher maturation of organoids at day 25. Moreover, we observed gene and TF expression patterns similar to the predictions based on the multi-modal analysis (Fig.4D and Supp. Fig. S13D-E;S16;S15). This included the TFs PAX3, RFX4 and ZIC2, which were predicted by PHLOWER to control the off-target neuronal lineage. Altogether, these results indicate that the Xenium panel can identify major differentiation branches in kidney organoids in a spatial context and will aid in understanding cell differentiation differences based on signaling events originating from cellular neighborhoods and spatial niches.

**Figure 4.**
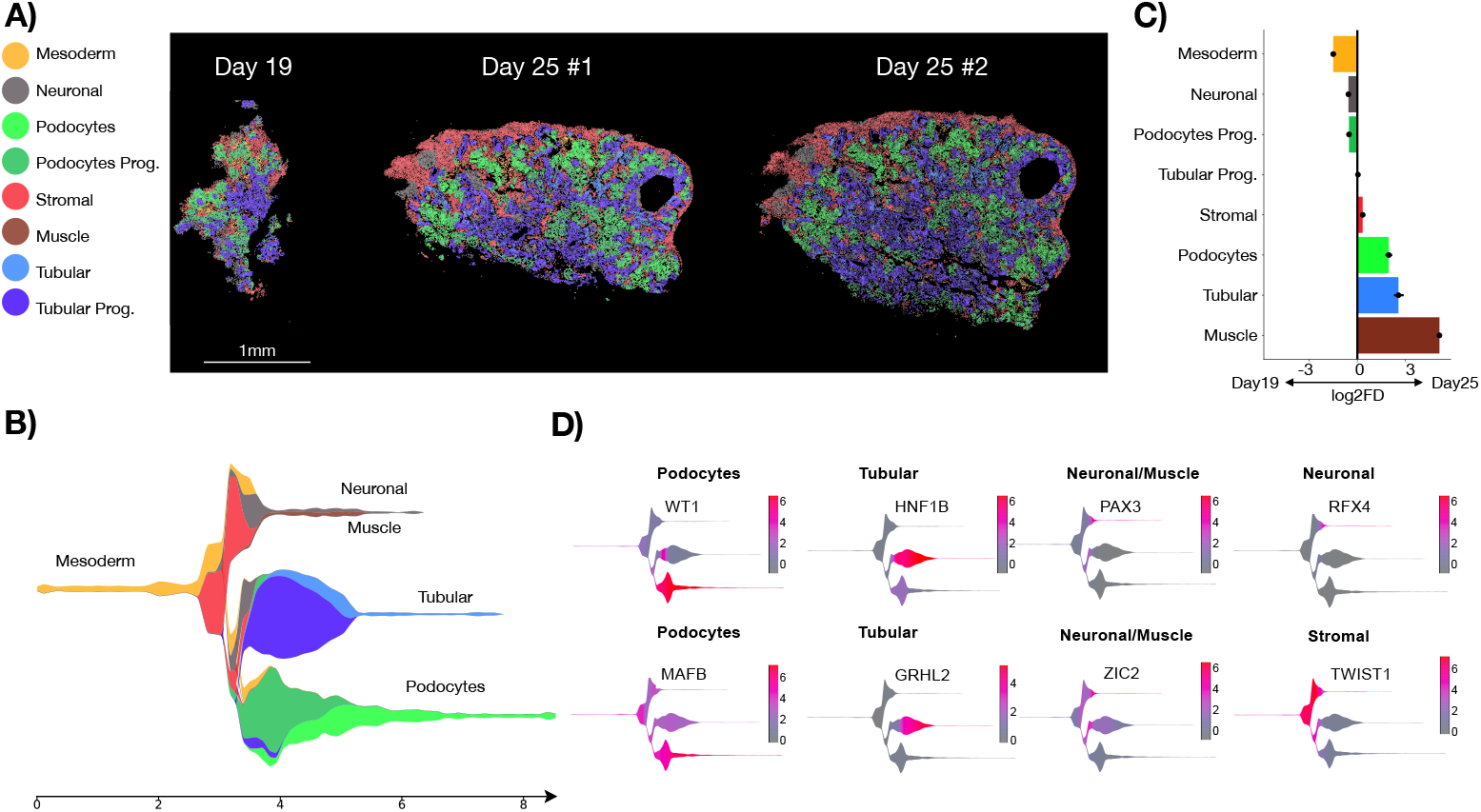
Spatial Profiling of Kidney Organoids. **A)** We show Xenium based spatial profiling of kidney organoids differentiation at day 19 and day 25 (section #1 and #2). Colors represent distinct cell populations. **B)** PHLOWER estimated differentiation tree on day 19 & day 25 organoids. **C)** Single cell proportion test comparing cell abundances in day 25 vs day 19. Error bars are based on confidence intervals for the magnitude difference returned via bootstrapping^60^. **D)** TFs controlling the differentiation of the kidney organoid branches.

To validate PHLOWER predictions, we performed multiplex siRNA knockdown experiments of the identified neuronal lineage defining TFs PAX3, RFX4 and ZIC2 in kidney organoids in iPSC derived kidney organoids during the differentiation process from day 19 onwards. Knock-down experiments led to a reduction of 25-30% in PAX3, RFX4 and ZIC2 mRNA expression (Supp. Fig. S14). Xenium spatial profiling of scrambled siRNA as control condition (scrambled; 2x section) and siRNA treated organoids against PAX3, RFX4 and ZIC2 detected the same cell types as non-treated day 25 organoids (Fig. 5A). When contrasting scrambled siRNA vs siRNA, we observed a significant increase of tubular cells and podocyte progenitors and a significant decrease of muscle, stromal and neuronal cells (Fig. 5B). Immunofluorescence staining further confirmed the reduction of stromal cells including off target neuronal cells in siRNA condition, as reflected by the significant decrease in interstitial vimentin (Fig. 5C-D). We observed a significant increase in podocyte nephrin protein expression and 50% more tubular E-cadherin protein expression, suggesting improved podocyte (progenitor) and tubular characteristics as a result of diminished off-target cells (Fig. 5E). Altogether, these experiments supported the that gene silencing of PAX3, RFX4 and ZIC2 known as neuronal lineage markers and considered as an off-target population, led to a small increase in tubular cell proportion and enhanced podocyte development as shown by increased nephrin and a higher cell proportion of podocyte progenitors.

**Figure 5.**
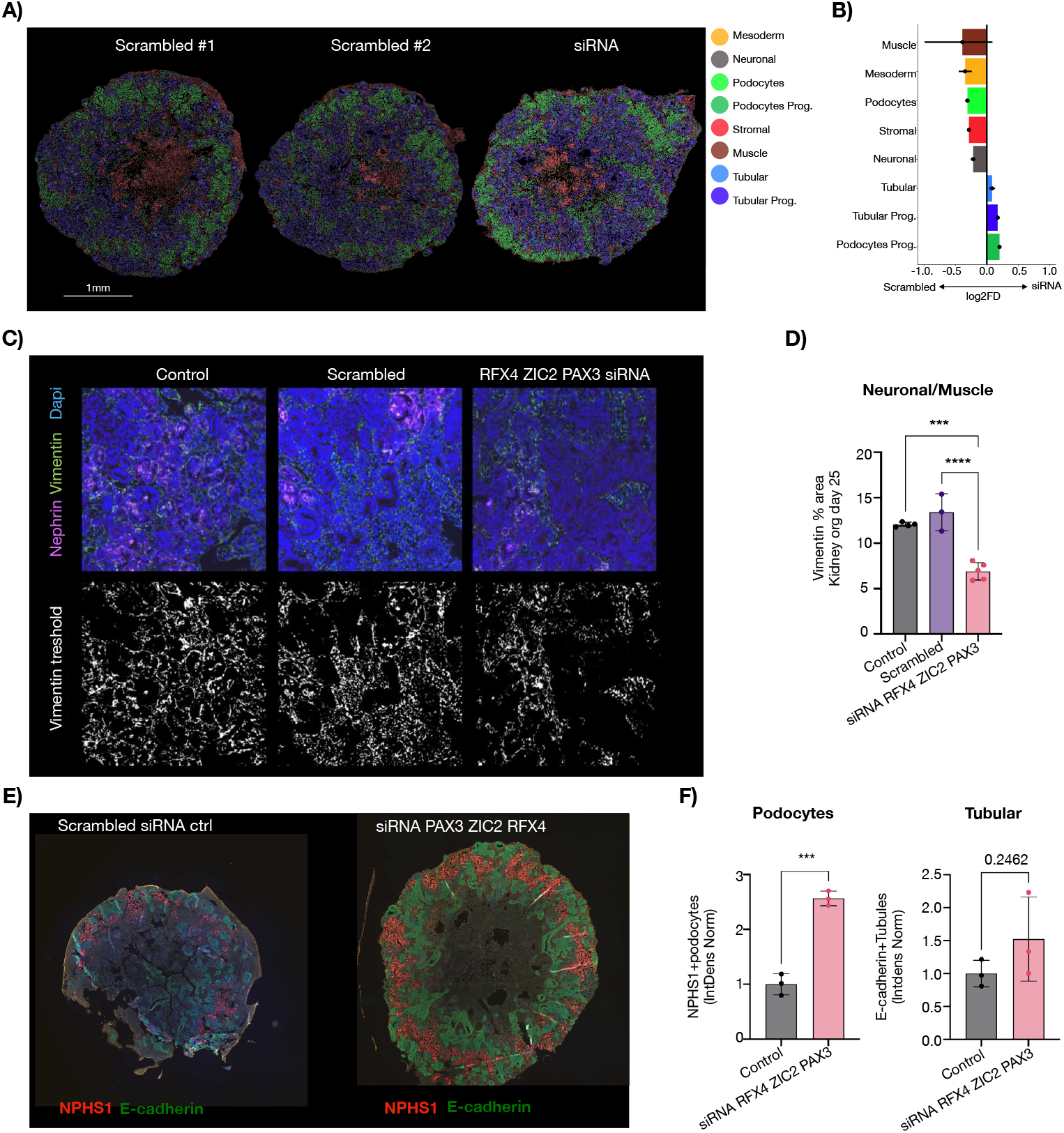
Spatial profiling of kidney organoids after siRNA knockdown of PAX3, RFX4 and ZIC2. **A)** We show Xenium based spatial profiling of kidney organoids at Day 25 after treatment with scrambled siRNA control (scrambled) and siRNA multiplex against the TFs PAX3, RFX4 and ZIC2 (siRNA). **B)** Single cell proportion test comparing cell abundances in siRNA and scrambled siRNA kidney organoids. Error bars are based on confidence intervals for the magnitude difference returned via bootstrapping^60^. **C)** Immunoflurecence staining of control. scrambled siRNA and siRNA treated kidney organoids showing nephrin (magenta), vimentin (green), and dapi (blue). **D)** Quantification of the neuronal marker vimentin^+^, NPHS1^-^ of images of (C) using one-way ANOVA analysis followed by Tukey post-test with bars representing mean ± sd; asterisks indicate P-values of <0.001(***), <0.0001(****). **E)** Silencing of RFX4, ZIX3 and PAX3 increased nephrin (red) expression in podocytes and increased e-cadherin (green) expression in tubular cells by 50%, though not significant. **F)** Podocyte and tubule quantification using unpaired *t*-test with bars representing mean ± sd; asterisks indicate P-values of <0.001(***).

## 3 Discussion

Trajectory analysis is paramount in the analysis of cells undergoing cell differentiation. Despite a wealthy literature of computational methods^8^, most methods have been only applied or tested in small cell differentiation trees with few branches. PHLOWER explores the harmonic component of the Hodge Laplacian decomposition, which allows to consider the interaction between edges, nodes, and triangles of a simplicial complex in order to estimate embedding on edges and trajectories levels. By exploring this higher-order representation of the data explicitly it allows the detection of complex and large cell differentiation trees, as supported by our benchmarking analysis on both simulated and real data sets.

Multimodal single cell data, which allows the parallel measurement of transcriptome and open chromatin data, can provide rich information on the relationship between regulatory and transcription function of cells^3^. We display the power of PHLOWER by analysing a kidney organoid differentiation course, where PHLOWER was able to detect major cell lineages, as well as TFs with branch specific gene expression and TF activity. We leveraged gene markers from PHLOWER predicted lineages to delineate a gene panel for uncovering the architecture of kidney organoids using in-situ sequencing spatial profiling. A particular important aspect of organoid growth is to improve protocols via the modulation of particular lineages^46,47,61^. Undesired off-target populations have been described to hamper kidney organoid development^46,47,61^. Targeting these populations using genetic or pharmacological perturbations led to improved organoid development and maturation. Here, we used siRNA targeting TFs controlling neuronal cells, as identified by PHLOWER, to reduce the abundance of these off-target cell types in kidney organoids and improved tubular cell formation. This reinforces the ability of PHLOWER to uncover regulators in complex cell differentiation systems.

An approach to characterize topological features in a data set is persistent homology^62^. Persistent homology can support PHLOWER by determining thresholds to build the triangulated SC on which we then compute the harmonic generators (Supp. Fig. S19). We also observed that persistence homology can be used to characterize the number of holes in the SC used by PHLOWER (Supp. Fig S20). However, distinctly from the harmonic components of the HL, persistent homology does not give unique generators that quantify the relation of *all* simplices (edges and nodes) to the holes ^63^. The combination of these approaches represent an interesting topic of further research.

Hodge Laplacians have been previously explored in other molecular problems^64^. HL of vector fields have been performed in RNA velocity estimates, which allowed visualization of curl-free, divergent-free and harmonic components^65^. This work also suggests that the harmonic component captured the overall direction of differentiation similarly as explored in PHLOWER. However, the previous work focused only on the visualization of the RNA velocity fields, and could not be used to infer the trees or to allocate cells along these trees as performed by PHLOWER. Another interesting application area of HL are three dimensional molecular data, as the prediction of protein ligands^66^. In such geometric data, higher order structures such as tetrahedrons (3-simplices) are also considered, which can reveal geometrical structures (2-loops) such as hulls or voids. This goes beyond the problem approaches here, which only require the identification 1-loops (holes). This is an interesting venue on research on the delineation of tissue structures from 3-dimensional spatial transcriptomics data^67^.

## 4 Methods

### 4.1 Rationale

PHLOWER uses the Hodge Laplacian (HL) and the harmonic component of the associated Hodge decomposition to obtain embeddings of cell differentiation trajectories. For this, PHLOWER represents single cell data as a simplicial complex consisting of nodes (0-simplices), edges (1-simplices) and triangles (2-simplices). Next, a first-order Hodge Laplacian is used to describe the simplicial complex. The first-order HL describes how edges relate to each other via nodes (so-called lower-adjacency) and triangular faces (upper-adjacency). Importantly, the decomposition of the HL can be used to decompose flows and trajectories on the edges into gradient-free, curl-free, and harmonic components, akin to the classical Helmholtz decomposition of vector fields known from vector calculus. While we focus here on “discrete” Hodge Laplacians based on simplicial complexes, there are guarantees that the spectral behavior of these converges to the Hodge Laplacian on manifolds in the limit if weighted accordingly^20,21^.

Of particular interest in our context here are the spectral embeddings associated to the so-called harmonic eigenvectors of the HL^11,18^, as these are associated to “holes” in the underlying space. Just like the eigenvectors of the graph Laplacian can be used to define a spectral embedding of the nodes in a graph, the eigenvectors of the Hodge-Laplacian can be used to provide an embedding of the edges. As edges correspond to cell differentiation events, this enables PHLOWER to represent cell differentiation events and cell differentiation pathways (sequences of edges) in a direct way. See Supp. Fig. S21 for a graphical contrast of the L0 and L1 Laplacian decompositions.

### 4.2 Overview of PHLOWER

PHLOWER receives as input a set of matrices representing a single cell sequencing modality that is:

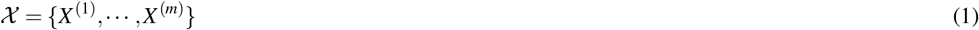

where *X* ^(*i*)^ ∈ ℝ^*n*×*s*(*i*)^ represents the data of a particular single cell modality, *n* represents the number of cells, and *s*^(*i*)^ represents the number of features in modality *i*. Here, we assume that cells match across modalities. PHLOWER has been evaluated on multimodal single cell (ATAC and RNA) sequencing or unimodal scRNA-seq (*m* = 1). First, PHLOWER constructs a graph representation *G* of the single cell matrix by estimating a joint embedding^9^ followed by kernel representation, graph construction and pseudo-time estimation as done in the literature^22^. Next, it builds a simplicial complex (SC). For this, it first uses the Delaunay triangulation procedure between nodes (cells) to obtain the set of 2-cells (triangles). Next, it uses pseudo-time information from the graph to find terminal differentiated cells and root cells. Finally, it connects terminal (differentiated) cells to root (progenitors) cells with edges and triangles to create holes corresponding to cell trajectories to obtain the final SC. Thus, PHLOWER created different cell differentiation paths measurable by topology/geometry, which we will exploit in the next steps.

PHLOWER next compute the harmonic eigenvectors of the normalized first order Hodge Laplacian^11^ associated to the SC. Eigenvectors with zero eigenvalues (harmonic eigenvectors) delineate holes in the SC, which are associated with cell differentiation trajectories. Lastly, PHLOWER sample trajectories from the SC which represent edge flows. Taking the dot product of individual edge flows with the harmonic eigenvectors creates a trajectory embedding, where each point represents a particular trajectory. Clustering analysis on this space allows PHLOWER to find major trajectory groups. Moreover, PHLOWER builds a cumulative trajectory space, which is used to recover the differentiation trees. This can be visualised as stream plots^5^. PHLOWER outputs a tree structure, the association of each cell within a branch and pseudo-time estimates (Supp. Fig. S1). It also detects branch specific regulators by detection of TFs similar as in^48^ with: (1) branch specific expression patterns and (2) similar gene expression and TF activity^49^ along the cell differentiation.

### 4.3 Single Cell Graph Representation

As a first step, PHLOWER estimates a single cell graph to represent the data using a procedure similar to DDHodge^22^. This procedure takes as input a low-dimensional embedding *X*^*l*^, which can be provided by MOJITOO^9^ for multi-modal data; or by principal component analysis (PCA) for scRNA-seq.

#### 4.3.1 Diffusion Map

Given a common joint cell embedding *X*^*l*^, we represent the data using diffusion maps^12^. For this, we first estimate a Gaussian kernel *W* :

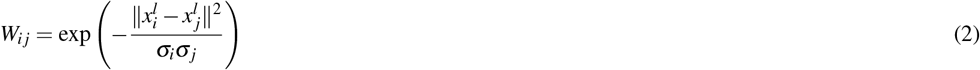

where 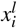 is the representation of sample (cell) *i* in the embedding *X*^*l*^ and *σ*_*i*_ is the local scaling parameter. This is estimated with the distance to the *n*-th nearest neighbor from 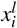 as in^68^.

The Graph Laplacian *L*_0_ can be defined as:

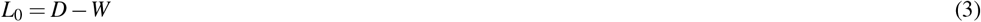

where *D* is a diagonal matrix with *i*-th entry *d*_*ii*_ = ∑ _*j*_ *W*_*i j*_. We now consider a random walk process on the graph described by the above graph Laplacian with

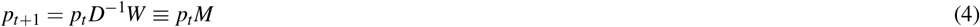

where *p*_*t*_ ∈ ℝ^*n*^ is the normalized probability vector at time *t, M* is the transition matrix of the random walk. Next, we perform an eigen-decomposition on the symmetric form of *M* such that:

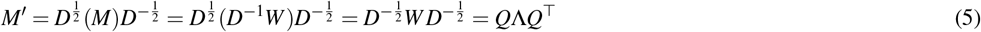

Taking advantage of the above eigendecomposition, we can effectively calculate *M*^*s*^ (see^12^), such that:

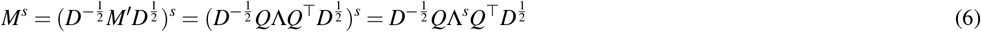

where the columns of 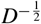 *Q* include the right eigenvectors of *M* and the rows of 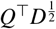 binclude the left eigen vectors. The diagonal matrix Λ contains the corresponding eigenvalues. Finally, we can estimate pseudotime *u* (or potential) at time *t* = *s* as:

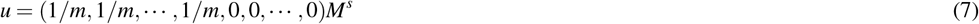

where *m* is the number of start cells, and *s* is the step of the diffusion process.

### 4.4 Graph-based pseudotime estimation

To improve the pseudotime estimates, we use a procedure similar to^22^, which smooths pseudo-time estimates by considering the graph connectivity. Let us denote the fully connected graph by *G*^*F*^ = (𝒱, ℰ ^*F*^) and the associated pruned *k*-nearest neighbor graph by *G*^*P*^ = (𝒱, ℰ ^*F*^). Next, we prune this fully-connected graph by considering only the *k* nearest neighbors. This provides a graph *G*^(*knn*)^ = (𝒱, ℰ), where 𝒱 is the set of vertices and ℰ is the set of edges. We represent the two graphs as an incidence matrices **B**_**1**_^*F*^ and **B**_**1**_^*P*^. For X ∈ {*F, P*}, vertex *v* _*j*_ ∈ 𝒱 and edge *e*_*i*_ ∈ ℰ ^*X*^, we have:

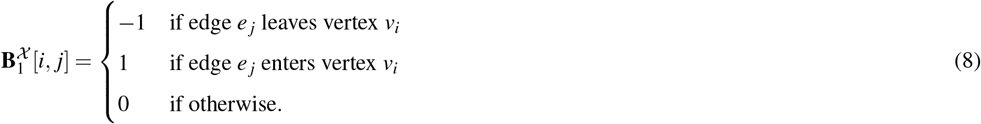

We now define initial edge weights as gradient values of the pseudo-time estimates of Eq. 7 on the fully-connected graph, e.g.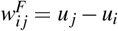. We want to estimate pseudo-times of the pruned *k*-nn graph by using the pseudo-time estimates of the full graph. For this, we get a first estimate of the gradients of the truncated graph (*w*^*P*^) by minimizing:

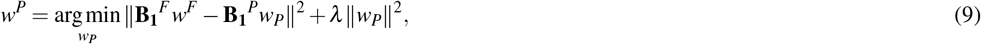

where *λ* is the regularization parameter. Next, we update the potential of the vertices

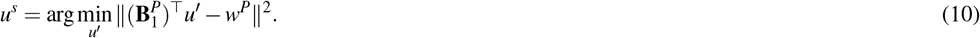

This allows us to estimate an updated gradients 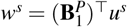.

With the graph *G*^(*knn*)^ = *G*^*P*^ with edge weights *w*^*s*^ and pseudotime estimates *u*^*s*^, we estimate a graph embedding with the stress majorization layout algorithm^15,69^. Examples of the graph embeddings and pseudo-time estimates can be seen in Supp. Fig. S1A.

### 4.5 Hodge Laplacian Decomposition on a Single Cell Simplicial Complex

Given *G*^(*knn*)^ and pseudo-time estimate *u*^*s*^, the next steps are the creation of a simplicial complex and estimation of the Hodge decomposition.

#### 4.5.1 Triangulation of the single cell graph

A typical cell differentiation graph has tree-like structures. The harmonic eigenvectors of the Hodge Laplacian are able to characterize distinct types of topological structures (connected components, holes, voids), but not trees or branches. PHLOWER resorts to a trick, i.e. it uses pseudo-time estimates to find end-state cells and adds dummy/artificial edges from end-state to start-state cells. This creates holes in the data, one for each trajectory group in the tree. For this, we obtain *m* (default 5) cells with highest and lowest pseudotime. We include edges from terminal (high pseudotime) to start (low pseudo-time) vertices. Next, we use a Delaunay triangulation^70^ on the graph embedding to create an hole-free set of triangles by first ignoring the dummy edges. Next, we remove edges connecting distant vertices, i.e. we estimate the distribution of distance of connected vertices and obtain the value associated with the 75% quantile (*Q*_75_) and we remove all edges connecting vertices, which distance is higher than 3 × *Q*_75_. We denote this as the **triangulated graph**. We further refine the dummy edges by performing a triangulation between each pair of terminal vertices and root vertices. This results in a **Simplicial Complex** *SC*= (𝒱^(*t*)^, ℰ^(*t*)^, 𝒯^(*t*)^), where 𝒱^(*t*)^ is the vertices set, ℰ^(*t*)^ is the edge set and 𝒯^(*t*)^ is the set of triangles. An example of the simplex representation of a differentiation tree is provided in Supp. Fig. S1B.

Remark: The filtering of edges by distance to obtain the **triangulated graph** is related to persistent homology analysis^62^. To show this, we evaluated how the proposed filtering from the **triangulated graph** compares to persistent homology analysis of connected components for the mouse embryonic data and simulated data (Supp. Fig. S19). This comparison suggests that using a filtering step such that we obtain one connected component plus an additional radius (1.2 of the minimum radius to obtain a single component) produces similar results as the previously described filtering of edges. The user can use either approach now in PHLOWER.

#### 4.5.2 Matrix Representation of a Simplicial Complex

Next, we represent the 2-dimensional simplicial complex *SC* = (𝒱^(*t*)^, ℰ^(*t*)^, 𝒯)^(*t*)^, as incidence matrices or boundary operators **B**_*k*_ which map *k*-simplices to (*k* − 1)-simplices. For example, the first boundary operator **B**_1_ captures the relation between vertices (0-simplices) and edges (1-simplices) and **B**_2_ captures the relation between edges (1-simplices) and triangles (2-simplices)^71^.

The boundary matrix on 1-simplices **B**_1_ is defined as in Eq. 8. An entry in **B**_2_ capturing the relationship between an oriented edge *e*_*i*_ ∈ ℰ^(*t*)^ and an oriented triangle Δ_*q*_ ∈ 𝒯 ^(*t*)^ can be defined as:

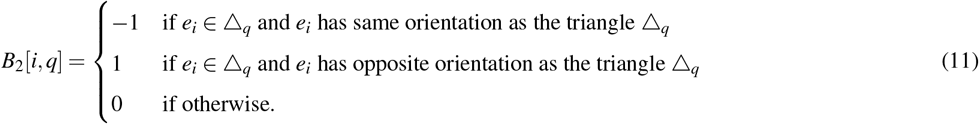

Check figure S22 for an example of a simplicial complex and its corresponding **B**_1_ and **B**_2_ matrices. There we can find several edges ([1, 2], [2, 3], [2, 4], [2, 5], [3, 4], [4, 5], [4, 6], [4, 7], [6, 7], [7, 8]) and 3 triangles ([2, 3, 4], [2, 4, 5] and [4, 6, 7]). Entry *B*_1_[1, 1] has value -1, as the first edge *e*_1_ = (1, 2) leaves vertex 1, while *B*_1_[2, 1] has value 1, as the first edge [1, 2] enters the vertex 2. Regarding **B**_2_, all entries related to the first edge *e*_1_ = (1, 2) are zero, as there is no triangle associated to it. *B*_2_[2, 1] is equal to one as the direction of *e*_2_ = (2, 3) fits the direction of the first triangle (Δ_1_ = (2, 3, 4)). *B*_2_[3, 1] is equal to -1 as the third edge (*e*_3_ = (2, 4)) has opposite direction to Δ_1_ = (2, 3, 4). We refer the reader to^11^ for an in-depth characterization of simplicial complexes. A similar rationale follows for **B**_2_. Note that for computational simplicity, the orientations of edges and triangles are set with a bookkeeping procedure^11^. That is, vertices are given numerical ids in order of creation; and edges and triangles are oriented towards vertices with higher id (see Sup. Fig. S22).

#### 4.5.3 Hodge Laplacian and Hodge decomposition

The Hodge Laplacian is a higher-order generalized form of the graph Laplacian. The *k*-th Hodge Laplacian is defined as:

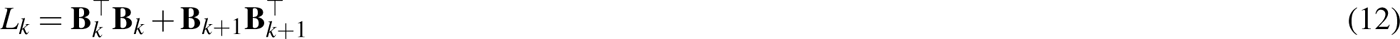

where **B**_*k*_ is the *k*-th boundary operator.

For *k* = 0, *L*_0_ is the same as the graph Laplacian introduced in Eq. 3.

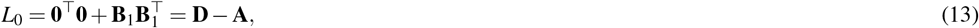

where **D** is the diagonal degree matrix of a graph and **A** is the adjacency matrix.

Here, we are interested in the first order Hodge Laplacian, i.e.:

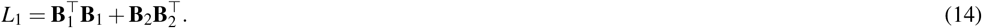

As in diffusion maps^23^, it is preferable to work with the normalized form of the Hodge Laplacian, as this provides a random walk process on the simplicial complex^11^:

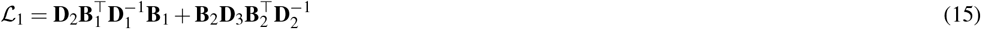

where **D**_2_ is the diagonal matrix of (adjusted) degrees of each edge, i.e. **D**_2_ = max(diag(|**B**_2_|**1**), **I**). **D**_1_ is the diagonal matrix of weighted degrees of the vertices according to the edge weights **D**_2_, and **D**_3_ = ^1^ **I**. Like the standard form of L_0_, decomposition hard. To address this, we construct the symmetric form of L_1_ with closely related eigenvectors and the same eigenvalues as following:

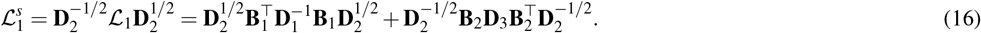

Next, we perform eigendecomposition on the symmetric form 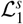 such that:

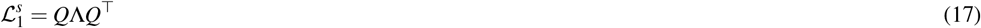

where columns of *Q* are the eigenvectors and the diagonal elements of the diagonal matrix Λ indicate the corresponding eigenvalues. Thus the decomposition of the normalized form L_1_ becomes:

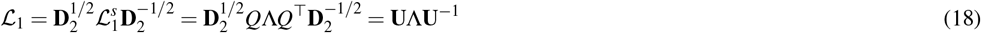

where 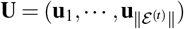 is the eigenvector matrix of 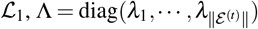 are the eigenvalues, and ‖ℰ^(*t*)^‖ is the number of edges in the *SC*. We assume the eigenvectors have been sorted by their corresponding increasing eigenvalues such that 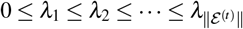. We denote.

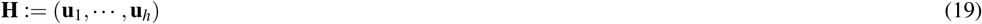

to be the matrix containing all the harmonic eigenvectors associated to ℒ_1_, i.e. all of the eigenvectors corresponding to the 0 eigenvalues, where 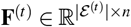.

For example, for the embryonic mouse data, we observe that the Hodge Laplacian spectra has two zero eigenvalues up to numerical inaccuracies, which define two harmonic eigenvectors. If we plot the harmonic eigenvector values on vertices, we observe that they highlight the two major branches of the triangulated graph (Supp. Fig. S1C). This is related to the spectral clustering algorithm, where eigenvectors with zero eigenvalues are associated with disconnected components (or clusters) in a graph (Supp. Fig. S21).

### 4.6 Trajectory Embedding and Tree Inference

#### 4.6.1 Trajectory Sampling and Embedding

To generate trajectories, we sample paths (or edge flows) in the *SC* by following edges with positive divergence (or increasing pseudotime). Due to the sparsity of the *SC*, we sample paths in *G*^(*knn*)^. For this, we perform a preference random walk on graph *G*^(*knn*)^. We choose a random starting point from the vertices (cells) with the *m* lowest pseudo-time values. We choose the next vertex randomly by considering the divergence values (*w*^*s*^). Only positive divergences (increase in pseudo-time) are considered. We stop when no further positive potential is available. Next, we return to the *SC*, estimate shortest paths between vertices in case sampled edges (from *G*^(*knn*)^) are not present in the *SC*. We define the embedding **f** ∈ ℝ|ℰ^(*t*)^| of a path on *SC* into the edge flow space as:

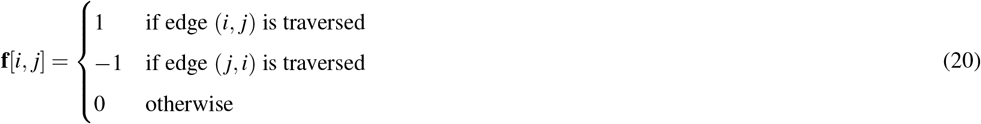

The random walk is repeated *n* times. This provides us with a path matrix **F**(*t*) ∈ ℝ|*ℰ* ^(*t*)^|×*n*, where |ℰ^(*t*)^| is the number of edges in the *SC*. See Supp. Fig. S1D for examples of sampled trajectories or edge flows. We next project these paths **F**(*t*) onto harmonic space to estimate a trajectory embedding:

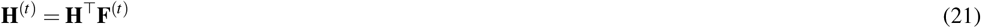

where **H** ∈ ℝ^*h*×*n*^ are the harmonic eigenvector defined in Eq. 19. PHLOWER next performs clustering on the trajectory embedding **H**(*t*) with DBSCAN^72^ to group the paths into major differentiation trajectories. For the mouse embryonic cell data, we observe that the trajectory embedding (or trajectory map) reveals two clusters S1E (left) associated with the neuronal and myocyte differentiation.

#### 4.6.2 Cumulative trajectory embedding and tree inference

The path representation presented in Eq. 20 does not keep the time step of a edge visit. Thus we also define a traversed edge flow (traversed path) matrix 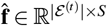 to record the edge visits for each time step individually, i.e.:

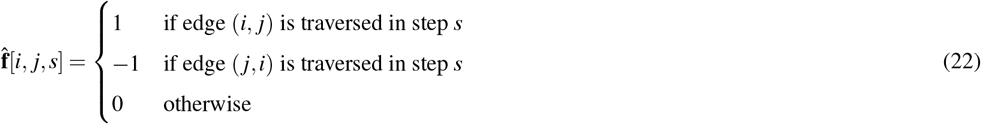

where *S* is the length of the trajectory, 1 ≤ *s* ≤ *S* is the number of the step and |ℰ(*t*)| is the number of edges in the graph *SC*. As we have *n* trajectories, we will have *n* traversed edge flow matrices 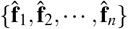.

We make use of cumulative trajectory embedding to represent paths and to detect major trajectories groups and branching points. For a path 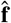, we can obtain a point associated with every step *s* in this cumulative (harmonic) trajectory embedding space as:

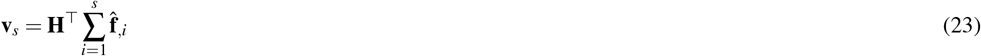

where 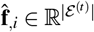 is a signed indicator vector for the edge traversed in step *i, v*_*s*_ ∈ ℝ *h* is the cumulative trajectory embedding of path 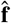 in step *s* in harmonic coordinates and *h* is the number of harmonic eigenvectors with zero eigenvalues. This is computed for all 1 ≤ *s* ≤ *S*, which defines a vector **v** = {**v**_1_, 00B7 · ·, **v**_*S*_} for every path. Intuitively, the entries of **v** describe a trajectory in the harmonic edge flow space that starts at 0 and ends in the harmonic trajectory embedding of the entire path.

As observed in Supp. Fig. S1E, these vectors are low-dimensional representations of paths in the cumulative trajectory embedding. By coloring paths from distinct groups with distinct colors, we can recognize branching point events, branches shared by trajectory groups and terminal branches. Note also that if we consider only the final entry *v*_*S*_ for every path, we obtain the same result as in the previously described trajectory embedding.

Recall that we have performed the DBSCAN clustering method to cluster the *n* paths into *m* groups {*g*_1_, *g*_2_, · · · *g*_*m*_} on the trajectory embedding defined in Eq. 21. PHLOWER next uses a procedure schematized in Supp. Fig. S23 to find the differentiation tree structure. First, it estimates pseudo-time values for every edge, i.e. the average pseudo-time *u*^*s*^ from vertices associated to the edge. It next bins all edges by considering their pseudo-time, i.e. it selects the trajectory group with the lowest pseudo-time and splits its edges in *p* bins (Supp. Fig. S23 B). The same range of pseudo-time is used to bin all trajectory groups and bins are indexed in increasing pseudo-time (Supp. Fig. S23 C). After binning of edges for each group, PHLOWER next finds the branching points for all group pairs by comparing the distance of edges within the bin vs. the distance of edges between the bins for a given bin index *i*.

More formally, for groups *g*_*i*_ and *g* _*j*_ and bin *k*, their corresponding edges in cumulative space are defined as set

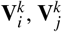. We then estimate the average edge coordinate per bin to serve as backbones for every group, i.e.:

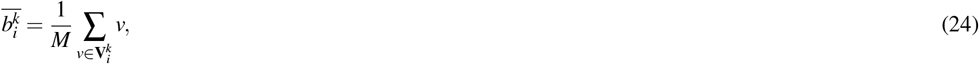

where 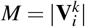 is the number of edges in the bin. We also consider the average distance between edges in a bin to have an estimate of the compactness of edges in a bin and trajectory, i.e.:

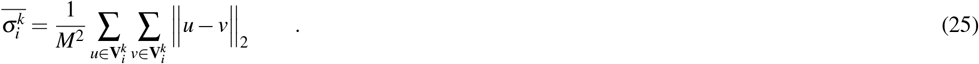

Finally, we calculate the distance between two groups *g*_*i*_ and *g* _*j*_ in time bin *k*, such that:

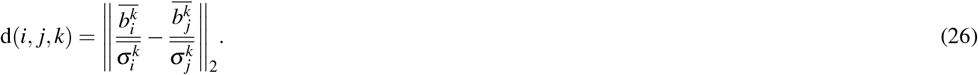

For every pair of groups, PHLOWER finds a unique branching point by trasversing bins in decreasing order and finding the first bin such that d(*i, j, k*) *< σ* (as default 1).

This is repeated for all pairs of groups. The tree is finally built in a bottom up manner. PHLOWER first considers branching points with highest index and builds a sub-tree by merging the two trajectory groups at hand. This is repeated until all branching points are considered (Supp. Fig. S23D-E). PHLOWER finally defines the leaves and the root by finding more extreme points, edges with lowest and highest pseudo-time in a trajectory group. These are the so called milestones (branching points, root and end points) needed for evaluation by Dynverse^8^. Trajectories are redefined as branches, i.e. part of the trajectories between two milestones. We also allocate cells (vertices) to these branches. For this, we consider the location of all edges associated with a vertex and use the mean value in the cumulative trajectory embedding space. This is used to allocate cells to branches and to find the distance between respective milestones. Moreover, we provide this information for STREAM^5^ for visualization as an STREAM tree.

#### 4.6.3 Regulator and Marker Selection

We make use of the cumulative trajectory space and statistical tests to find markers and regulators associated to trajectories. Two compare two final branches, PHLOWER selects all cells associated within particular areas of the branches, i.e. highest 50% of the bins. Note that every cell can be visited by several edges. To consider this important information, we multiply expression count vector of each cell by the number of visits. We then perform a statistical test (default is the *t*−test from scanpy^73^). In case we are interested in comparing sub-trees with several branches, PHLOWER first merges the sub-tree as a single branch, before the selection of the bin.

In the case of multimodal data (RNA and ATAC), PHLOWER also has a module to find regulators. First, we estimate a TF activity score per cell using chromVar^49^. We then use the previously described test to find branch specific regulators. Since TF activity cannot be discerned from TFs with similar motifs, we only consider genes, which TF activity is similar to the expression pattern at the selected branch as in^48^. We first get the average expression/TF activity of cells of bins around the branch of interest. We next smooth the gene expression using convolution (numpy.convolve)). We then estimate the correlation between gene expression and TF activity and use this to sort branch specific regulators.

### 4.7 Materials

#### 4.7.1 Synthetic data sets

To test the power of PHLOWER in the detection of multiple branches trees, we generate 10 simulated data sets using diffusion-limited aggregation trees (DLA tree)^23^ as proposed in PHATE^14^. For this, we use a Gaussian noise parameter of 5 and vary the parameter n_branch, which controls the number of branches in the trees, from 3 to 12. This generated trees with 5 to 18 branches. For every data, we generate 3000 points and 100 features. Of note, we use the same data as evaluated in PHATE as data with 10 branches. Next, we run PHATE to visualize the branches, with which we can future construct ground truth for dynverse benchmarking.

Next, we reformat this data to be used within the trajectory benchmarking framework dynverse^8^ by using dynwrap. In short, dynverse needs detailed information of the branches, branch points and association of cells within a branch. To do so, we need to find the branching points of the DLA tree. First, we perform PHATE on the DLA high dimensional data to get embedding with 2 dimensions. We only consider embeddings, where the tree structure is preserved. Next, we find the branching points by finding 2 nearest neighbors of two branches. With these branching points, we created the branch backbones needed by dynverse. We calculate the association of each data point to a branching point (milestone percentage) by calculating the euclidean distance between the points related to each branch.

#### 4.7.2 Real scRNA-seq data sets

To evaluate the performance of PHLOWER on real datasets, we selected 20 real datasets from the Dynverse benchmarking dataset, including 4 gold-standard and 16 silver-standard datasets with known ground-truth tree structures. Specifically, we only consider data with a single root and at least three branches, while excluding subsets that were repetitive of the Planaria-full_plass dataset. See Supp. Table S1. For this, we checked out the code from https://github.com/dynverse/dynbenchmark and ran the script to download all the necessary data.

### 4.8 Benchmarking Evaluation

Dynverse includes wrappers for several trajectory methods. We explore the following methods in our evaluation: PAGA tree^4^, monocle 3^25^, slingshot^6^, Slice^28^, TSCAN^31^, Slicer^32^, MST^33^ and Elpigraph^34^. We expand this by including a wrapper for STREAM^5^ and PHLOWER. We first calculate the distance of each cell to all branches, next assign the cell to the nearest branch and use the distance ratio to associate cells to milestones. Next, we use dyneval to measure quality metrics for the generated trajectories. As in^8^, we mainly focus on HIM(Hamming-Ipsen-Mikhailov distance) to measure the tree structure similarity; *cor*_*dist*_ to measure the correlation between the cell geodesic distances within predicted and true branches; *F*1_*branches*_ and *F*1_*milestones*_ to measure the accuracy of a cell assigned to the correct branches and the correct milestone (bifurcation points).

### 4.9 Kidney Organoids

#### 4.9.1 Ethical statement

Permission for the creation and use of iPSCs in this study was obtained from the ethical commission at RWTH Aachen University Hospital (approval number EK23-193).

##### Cell culture

Generation of human iPSC-353 line: Erythroblasts from a healthy volunteer were reprogrammed using the CytoTune^™^-iPS 2.0 Sendai Reprogramming Kit (Thermo Fisher) according to the manufacturer’s protocol. In short, erythroblasts were transduced using the CytoTune^™^-iPS 2.0 Sendai Reprogramming Kit and re-seeded on plates with inactivated MEFs to support the growth and pluripotency of iPSCs. Emerging iPSC colonies were picked, expanded and assessed for activation of stem cell markers to confirm pluripotency.

##### Generation of human induced pluripotent stem cell (iPSC) derived kidney organoids

Generation of human induced pluripotent stem cell (iPSC) derived kidney organoids iPSC were seeded using a density of 20,000 cells/cm2 on Geltrex (Thermo Fisher, Breda, the Netherlands) coated 6-well plates (Greiner Bio-one, Alphen aan de Rijn, the Netherlands). The differentiation protocol was based on our previous work^39^. In brief, differentiation towards ureteric bud-like and metanephric mesenchyme lineage was initiated using CHIR-99021 (10 *µ*M, BioTechne Ltd., Abingdon, United Kingdom) in STEM^™^diff APEL^™^2 medium (StemCell Technologies, Cologne, Germany) for 3 and 5 days, respectively. Next, the medium was replaced sequentially for APEL^™^2 supplemented with fibroblast growth factor 9 (FGF9), 200 ng/ml, R&D systems) and heparin (1 *µ*g/ml, Sigma-Aldrich, Zwijndrecht, Netherlands) up to day 7. On day 7, differentiated cells were trypsinized and mixed in a ratio of one part 3 days CHIR-differentiated cells and two parts 5 days CHIR-differentiated cells to stimulate crosstalk between both lineages in order to boost segmented nephrogenesis. To generate cell pellets, 300,000 cells per 1.5 ml tube were aliquoted from the cell mix and centrifuged 3 times at 300 rcf for 3 min changing position by 180^°^ per cycle. Cell pellets were plated on Costar Transwell filters (type 3450, Corning, Sigma-Aldrich), followed by a one-hour CHIR pulse (5 *µ*M) in APEL^™^2 medium added to the basolateral compartment. Next, medium was replaced for APEL^™^2 medium supplemented with FGF9 and heparin for additional 5 days and the entire 3D organoid culture was performed using the air-liquid interface. After 5 days of organoid culture, APEL^™^2 medium supplemented with human epidermal growth factor (hEGF, 10 ng/ml, Sigma-Aldrich). Medium was replaced every other day for an additional 13 days.

##### Silencing of ZIC2, RFX4 and PAX3 during stem cell-derived kidney organoid development

Kidney organoids (at least N=3 per condition per experiment, 2 independent experiments) were transfected using ON-TARGETplus^™^ SMARTpool siRNA’s ZIC2 (Catnr L-017505-00-0005), RFX4 (catnr L-013577-00-0005) and PAX3 (Catnr L-012399-00-0005, work concentration 25nM, Horizon Discovery, Cambridge, UK) and DharmaFECT Transfection reagent 1 (0.5% v/v) from day 7+5 (intermediate mesoderm stage) onwards until the end of the protocol until day 7+18. Organoids were refreshed every other day.

##### RNA extraction, cDNA synthesis and qPCR

RNA from organoids was extracted using the PureLink RNA mini kit (Thermo Fisher) according to the manufacturer’s protocol. RNA was stored at -80^°^*C* until further processing. cDNA synthesis was performed using 200 ng RNA as input, using the iScriptTM cDNA synthesis kit (Bio-rad) according to the manufacturer’s protocol. The mRNA expression was quantified by performing a semi-quantitative real-time PCR using PowerUp SYBR Mastermix (Applied Biosystems) and primers targeting human RFX4, ZIC2, PAX3 genes (hZIC2_For AAAGGACCCA-CACAGGGGAGA, hZIC2_Rev GACGTGCATGTGCTTCTTCCT hRFX4_For TGGGAAGAGCATGCATTGTG hRFX4_Rev TCTTTCAATCCAGCTCTCTGTGG hPAX3_For GCAGTATGGACAAAGTGCCT hPAX3_Rev CAGGGCCAGTTTTAGCTCCA. After correction with the corresponding human GAPD gene (GAPDH_For GAAGGTGAAGGTCGGAGTCA, GAPDH_Rev TGGACTCCACGACGTACTCA), gene expression levels were plotted as fold change compared to control. For primer sequences please refer to the key resources table.

##### Harvesting kidney organoids for 10x genomics NEXT GEM multi-ome pipeline

To dissect nephrogenic differentiation trajectories in kidney organoids using the multiome pipeline, organoids were harvested at different time points during culture. The following differentiation stages were processed: day 7 (cells harvested from the 2D cell layer, primitive streak - intermediate mesoderm stage), organoids day 7+5 (day12), day 7+12 (day19) and day 7+18 (day25). These kidney organoid stages represent nephrogenesis ranging from intermediate mesoderm (day 7+5) towards metanephric mesenchyme and ureteric bud-like lineages (day 7+12) that result into nephron-like structures embedded by a (progenitor) stromal compartment at the end of the differentiation protocol (day 7+18). Kidney organoids (N=4 per timepoint) were harvested on the respective dates (d7+5, d7+12, and d7+18) by cutting single organoids out of the transwell filter in a sterile flow hood using a scalpel. Organoids were washed with 5 ml PBS per filter at room temperature 3 times. Afterwards, single organoids were cut from the transwell filter with the organoids still being attached to the membrane, transferred to 1.7 ml cryovials (Greiner bio-one) and subsequently snap frozen and stored at -80^°^*C* until nuclei isolation.

##### Single nuclei isolation from kidney organoids

Snap frozen kidney organoids were thawed in PBS and crushed using a glass tube and douncer (Duran Wheaton Kimble Life Sciences, Wertheim/Main, Germany). After passing the single cell suspension through a 40 µm cell strainer (Greiner bio-one), the suspension was centrifuged at 4^°^*C* and 300xg for 5 min. Subsequently, the supernatant was discarded and the cell pellet was resuspended in Nuc101 cell lysis buffer (Thermo Fisher), supplemented with RNase and protease inhibitors (Recombinant RNase Inhibitor and Superase RNase Inhibitor, Thermo Fisher, and cOmplete Protease Inhibitor, Roche), incubated for 30 seconds and centrifuged at 4^°^*C* and 500xg for 5 min. After discarding the supernatant, the nuclei were carefully resuspended in PBS containing 1% (v/v) Ultra-Pure BSA (Invitrogen Ambion, Thermo Fisher) and Protector RNAse inhibitor (Sigma Aldrich). Single nuclei were counted using Trypan blue (Thermo Fisher) and prepared for 10x genomics ChromiumNextGEM Multiome pipeline v1 according to the manufacturers guidelines (https://cdn.10xgenomics.com/image/upload/v1666737555/ support-documents/CG000338_ChromiumNextGEM_Multiome_ATAC_GEX_User_Guide_RevF. pdf).

##### FFPE tissue of human iPSC-derived kidney organoids

iPSC-derived kidney organoids were cut from the Transwell^™^ filter and fixated in 4% (v/v) formalin on ice for 20 min. Fixed iPSC-derived kidney organoids were stripped of the filter membrane using a scalpel. Each single organoid was embedded using 2.25% (w/v) agarose gel (Thermo Fisher). After embedding for 5 min at 4°*C*, the iPSC-derived kidney organoids were transferred to embedding cassettes and paraffinized. After paraffinization, iPSC-derived kidney organoids were cut at a thickness of 4 *µ*m using a rotary microtome (Microm HM355 S, GMI) and mounted on FLEX IHC microscope slides (DAKO, Agilent Technologies) for immunofluorescence staining or Xenium array slides for spatial analysis.

##### Immunofluorescence staining

Slides were deparaffinized using series of xylol (2^×^) and 100% (v/v) ethanol (3^×^), followed by antigen retrieval by boiling slides in Tris-buffered EDTA (VWR Chemicals) for 20 minutes. Primary (1:100) and secondary (1:200) antibodies were diluted in PBS containing 1% (v/v) bovine serum albumin (BSA; Sigma-Aldrich). Primary antibodies (NPHS1, AF 4269-SP, RD Systems, Vimentin, ab92547, Abcam, E-cadherin, 610405, BD Biosciences) were incubated overnight at 4^°^*C*, and secondary antibodies were incubated at RT for 2 h (donkey anti-sheep IgG (H+L) alexa fluor 647 (ThermoFisher), donkey anti-rabbit IgG (H+L) alexa fluor 488 (ThermoFisher), donkey anti-mouse IgG (H+L) alexa fluor 488, DAPI (300nM, 4’,6-Diamidine-2’-phenylindole dihydrochloride, Merck). After each antibody incubation, slides were washed three times in PBS for 5 min. Slides were mounted using Fluoromount-G^®^ (Southern Biotech, SanBio). Images were captured using a Zeiss LSM 980 confocal microscope.

#### 4.9.2 Computational Analysis of Kidney Organoids

After read mapping using the cellranger-arc tool(version 1.0.1), we filter the low-quality cells using information both from scRNA and scATAC reads. We first import scRNA to Seurat^74^ to get the scRNA metrics for filtering. We next import the fragments into ArchR^75^ to get the scATAC metrics for filtering. With the information above, we retain cells with barcodes in both scRNA and scATAC count matrices. Next, we filter the cells using threshold nFeatureRNA > 400 & nCountRNA <400,000 &percent.mito > 5 and scATAC using thresholds in ArchR that minTSS = 6 & minFrags = 2500 & maxFrags = 1e+05. We next perform preprocessing to scRNA using Seurat, to do this, we first normalize the scRNA data by calling NormalizeData with the default parameters, next we find top variable features with parameter selection.method=‘vst’. Next, we scale the data by regressing out the cell cycle and mitochondrial effect. We next run PCA with 50 PCs. To remove the batch effects, we next run harmony^40^ to integrate the four samples. For scATAC data, we use ArchR to perform the preprocessing. We first create Arrow files using the filter threshold aforementioned by calling createArrowFiles, the tile matrices are created directly from the fragment files. Next, we add doublets score by calling addDoubletScores for each sample followed by function filterDoublets to remove doublets with default parameters. Next, we run addIterativeLSI on the tile matrices to add dimensional reductions. A batch correction is also called using Harmony^40^ with function addHarmony. To obtain a uniform dimension reduction for the downstream analysis, we next run MOJITOO^9^ to integrate the scRNA and scATAC data with default parameters.

MOJITOO embedding was used as input for PHLOWER. An important parameter is the indication of the root cells. For this, we performed a clustering analysis using Seurat^74^ with resolution=2.5 on the MOJITOO embedding space. The root cells were defined by clusters predominantly present in day 7 and with the expression of mesoderm markers (TBXT, MESP1, KDR5). PHLOWER found 28 dimensions with zero eigenvalues and clustering analysis of the trajectory space detected 16 groups. We removed 4 main trajectories outliers (less than 0.5% samples inner the cluster) and one trajectory, where cells had a low pseudo-time values (e.g. they did not differentiate). Finally, some clusters had the same end time points. We therefore keep the one with the largest number of trajectories. This resulted in the 9 final trajectories found by PHLOWER.

#### 4.9.3 Computational Analysis of Xenium Experiments of Kidney Organoids

To select markers, we used the phlower.tl.tree_mbranch_markers function to identify markers for each main branch (stromal, neuronal, tubular and podocytes), focusing on cells with highest pseudo-time according to PHLOWER (top 50%). Next, we further selected markers with expression specific to sub-branches (tubular, podocytes, stromal 1, stromal 2-4, neuron 1, and neuron 2-3). For this, we retained only genes with a padj < 0.05 and logFC > 1.5 for the main branch markers. Next, we only considered protein-coding genes according to Ensembl (Version GRCh38.104). We then scaled the logFC values of both main and sub-branches to a 0-1 range and calculated the mean normalized logFC for each branch and its sub-branches. We selected the top 12 genes for each sub-branch based on the mean normalized logFC. For TF selection, we used TF-Gene correlation along each sub-branch to identify 5 TFs per main branch using scMEGA analysis shown in Supp. Fig. S12. Additionally, we included literature based cell type markers reported in Supp. Fig. S10, which supported the annotation of the multiome single cell data. In total, we identified 100 genes for the Xenium experiment (see Supp. Table S2). The panel primers were designed with the Xenium Panel Designer tool provided by 10X Genomics.

For the data analysis, we used Xenium Explore3 to inspect each region and remove those that did not show differentiation and remove stromal cell patches around organoid edges. This process left us with two scrambled siRNA regions, one siRNA region, one day 19 region, and two day 25 regions. Note that multiple regions corresponds to distinct sections of the same organoid. Next, we used Space Ranger with default parameters and loaded each Xenium sample using Seurat (version 5.0.2) with the LoadXenium function and removed cells based on the organoid masks. We integrated all cells using Seurat’s merge function and filtered out cells with zero nCount_Xenium. We next applied SCTransform for data normalization. PCA for dimensionality reduction and Louvain cluster (50 PCs and resolution = 0.3); and estimated a UMAP for visualisation. Next, we annotated the clusters by examining the gene expression of known cell type markers. We excluded 5 clusters with 33,396 cells due to low nFeature_Xenium(<20) and nCount_Xenium (<60). This potentially represents cells, whose markers are not present in our panel.

For the day 19 and day 25 regions, we performed trajectory inference using PHLOWER using a subset of the cells (2000 per cluster). We used Mesoderm cells as the root for the trajectory inference and identified three end branches. Finally, we integrated PHLOWER with STREAM for visualization.

### 4.10 Pancreas and Neurogenesis scRNA-seq data

We apply PHLOWER and competing approaches to pancreatic endocrinogenesis data (3,696 cells)^36^ and the data on developing mouse hippocampus (neurogenesis - 18,213 cells)^37^, both of which were used in scVelo^38^. Specifically, we extract AnnData using the scVelo package via scvelo.datasets.pancreas and scvelo.datasets.dentategyrus_lamanno. For both datasets, we appliced the Scanpy pipeline including filtering, feature selection, normalization and PCA with default parameters. See tutorials for details on how PHLOWER was executed in these datasets.

For competing approaches, these methods were executed with Dynverse. Basically, we first use R to load the h5ad file with anndata::read_h5ad and create a SeuratObject from the count matrix. We then perform normalization and identify the top 2,000 variable features to obtain the log-normalized expression matrix. Next, we use dyn-wrap::wrap_expression to structure a dynverse dataset object with count and expression data. The start_id is set as the first “Ductal” and “nIPC” cell ID for Pancreas and Neurogenesis, and group_id is assigned based on cell types. The dataset object is then saved as an R object, enabling trajectory inference with dynwrap::infer_trajectory. Finally, we visualize the inferred trajectories using dynplot::plot_dimred.

## Code and data availability

Code, documentation and examples for running analysis of this manuscript are available on github https://github.com/CostaLab/phl and readthedocs https://phlower.readthedocs.io/en/latest/. Data objects with the kidney organoid (multiome and Xenium) as well as the benchmarking data have been deposited in zenodo.

## Acknowledgements

This project has been funded by the German Research Foundation (DFG) (3888802535, CRU344-417911533 and CRU5011 - 445703531) and the Consortia E:MED Fibromap and CureFib funded by the German Ministry of Education and Science (BMBF). This work was further supported by grants of the German Research Foundation (DFG: SFBTRR219 322900939, CRU344-4288578857858, CRU5011-445703531), by a grant of the European Research Council (ERC-COG 101043403), the Dutch Kidney Foundation (DKF), TASKFORCE EP1805 and the NWO VIDI 09150172010072. MTS and VG acknowledge funding from the German Research Foundation (DFG) within Research Training Group 2236 (UnRAVeL) and the European Union (ERC, HIGH-HOPeS, 101039827). Views and opinions expressed are however those of the author(s) only and do not necessarily reflect those of the European Union or the European Research Council Executive Agency. Neither the European Union nor the granting authority can be held responsible for them.

## 5 Supplement

**Figure S1.**
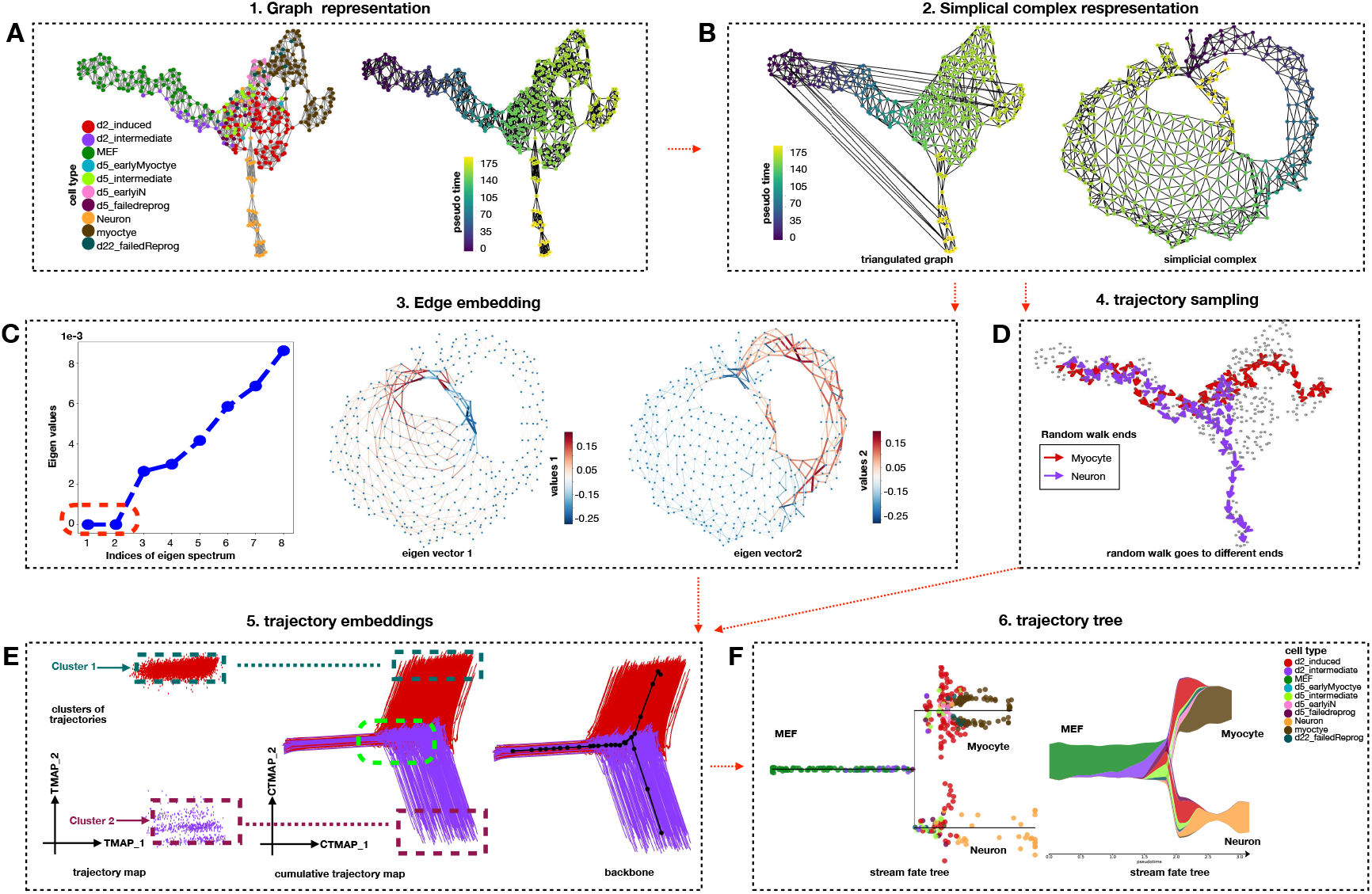
Detailed steps of PHLOWER on a single cell data with mouse embryonic fibroblasts towards neurons and myocytes. A) First, PHLOWER estimates a graph representation using kernels and stress majorization layout^15^. Colors correspond to cluster labels as defined in^16^. This information (labels) is not used by PHLOWER algorithms B) Next, a pseudo-time is estimated using the zero-order Laplacian and using MEF cluster as root. The definition of a root is the only supervision required in PHLOWER. Edges between cells are obtained via Delaunay triangulation (left). To create holes in the simplicial complex, PHLOWER performs a trick by including edges between high pseudo-time towards low pseudo time vertices (middle). If we re-run the graph layout, two holes are clearly visible. C) PHLOWER next computes the harmonic eigenvectors of the Hodge Laplacian of the simplicial complex. This indicates two harmonic eigenvectors, i.e. eigenvectors with zero eigenvalues. By plotting harmonic eigenvector’s values on the edges, we observe that the values of the first harmonic eigenvector discriminates edges in the neuronal branch vs. others, while the values of the second harmonic eigenvector discriminates the myocyte related cells from others. D) Next, PHLOWER generates trajectories by random walk on the simplicial complex representation to obtain edge-flow vectors. E) Finally, PHLOWER creates a trajectory embeddings (Eq. 21) by a dot product of the edge flows with the harmonic eigenvectors. We observe two clear clusters in the trajectory map (left), which are detected by providing this embedding as input for DBSCAN^72^. We can similarly create a cumulative trajectory map (Eq. 23) by creating edge flows of trajectories by considering only the first one, two, three and so on first differentiation events. PHLOWER uses this space to delineate the backbones of differentiation trajectories and to detect branching points events (in green, right panel). F) PHLOWER outputs a differentiation tree, pseudo-time values and an allocation of cells to positions in the branches. PHLOWER makes use of STREAM to visualise the final trajectories.

**Figure S2.**
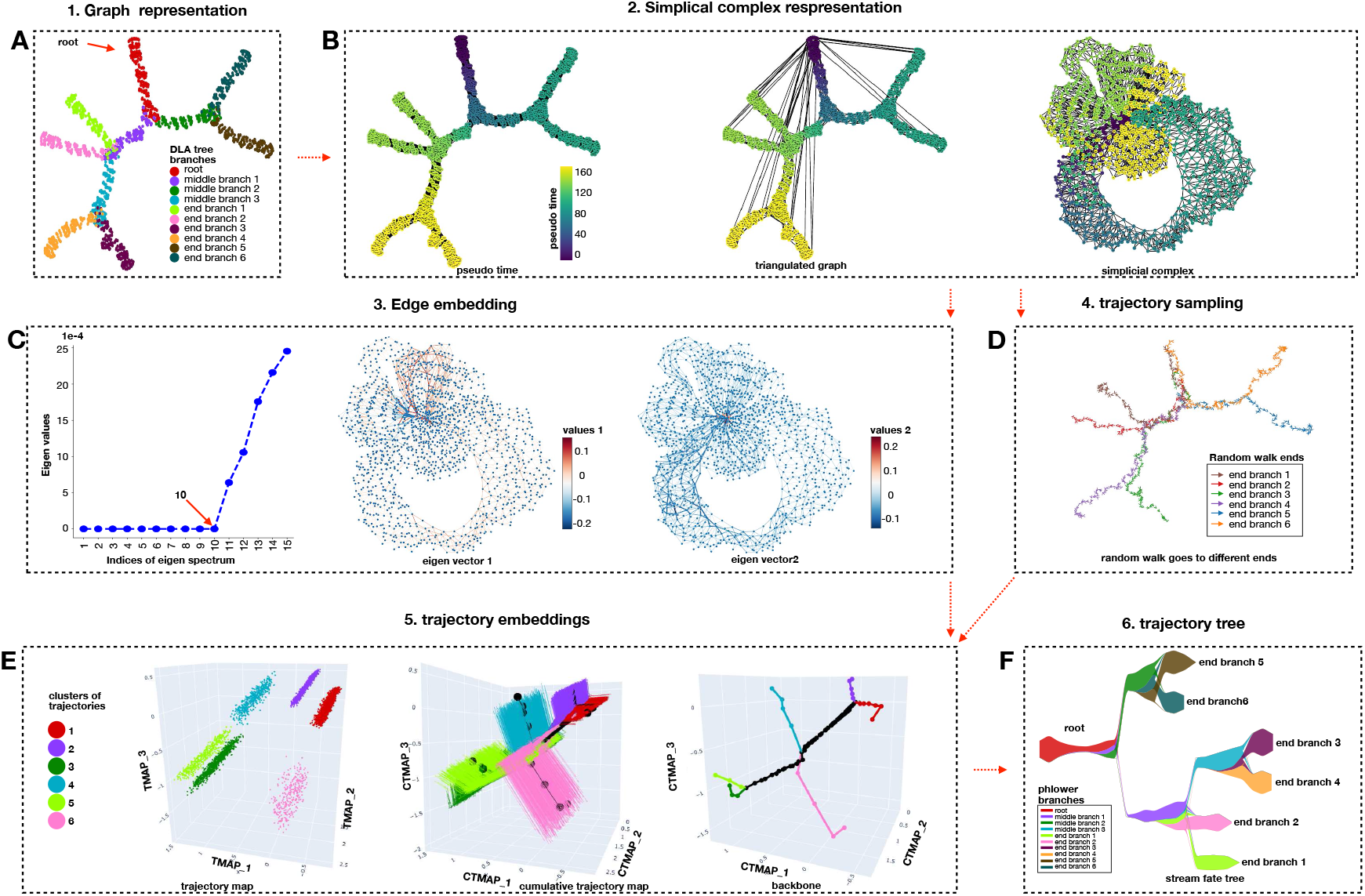
PHLOWER workflow for the simulated tree dataset with 10 total branches. Panels A-F show distinct PHLOWER estimates for the simulated data with 10 branches exactly as described in Supp. Fig. S1. Note that in this complex multi-branched data, the 2 dimensional representation of the graph layout of the simplicial complex (B) can not capture the data complexity and does not properly display all the holes. Clustering analysis in the trajectory embedding detects the six main trajectories present in the data (E) and recovers an almost perfect differentiation tree (F). Of note, the number of zero-eigenvalues (10) is higher than the number of main trajectories (6). This is potentially due to small (and noise associated) holes in the graph. This do not impact the precision of the clustering analysis to find the correct branches.

**Figure S3.**
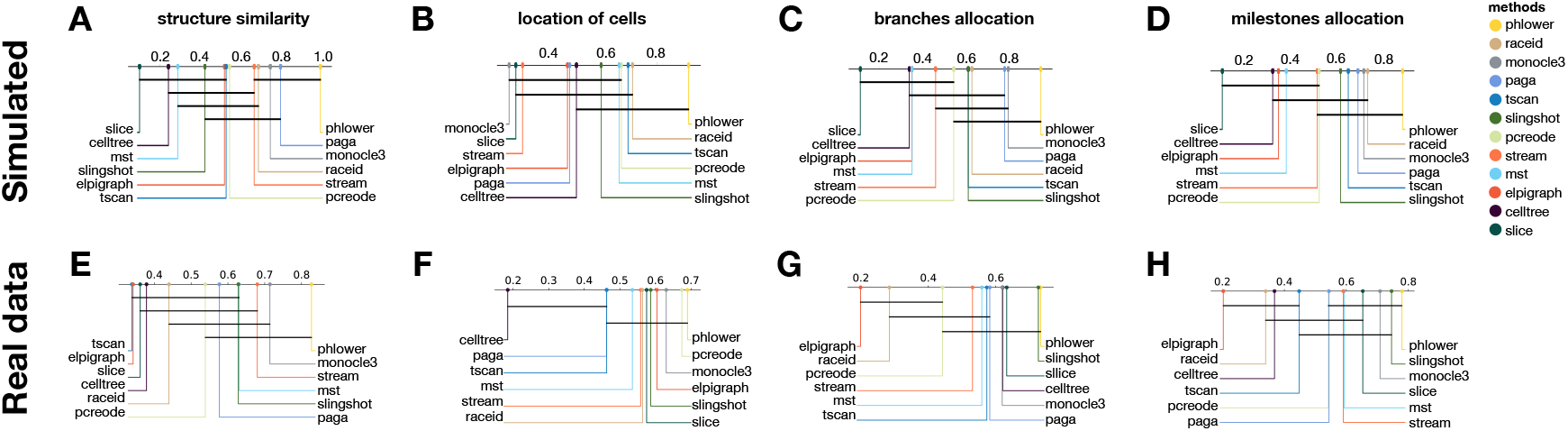
Ranking of Bechmarked methods: **A-D** We show the mean rankings of the methods based on the HIM, Correlation, F1 branches and F1 milestones for all evaluated methods in the simulate data set (based on results from Fig.2A-D). Higher ranking values indicate the best performer. Bars indicate methods with similar performance in accordance to the Friedman-Nemeniy test. E-H Same as A-D for the scRNA-seq data sets.

**Figure S4.**
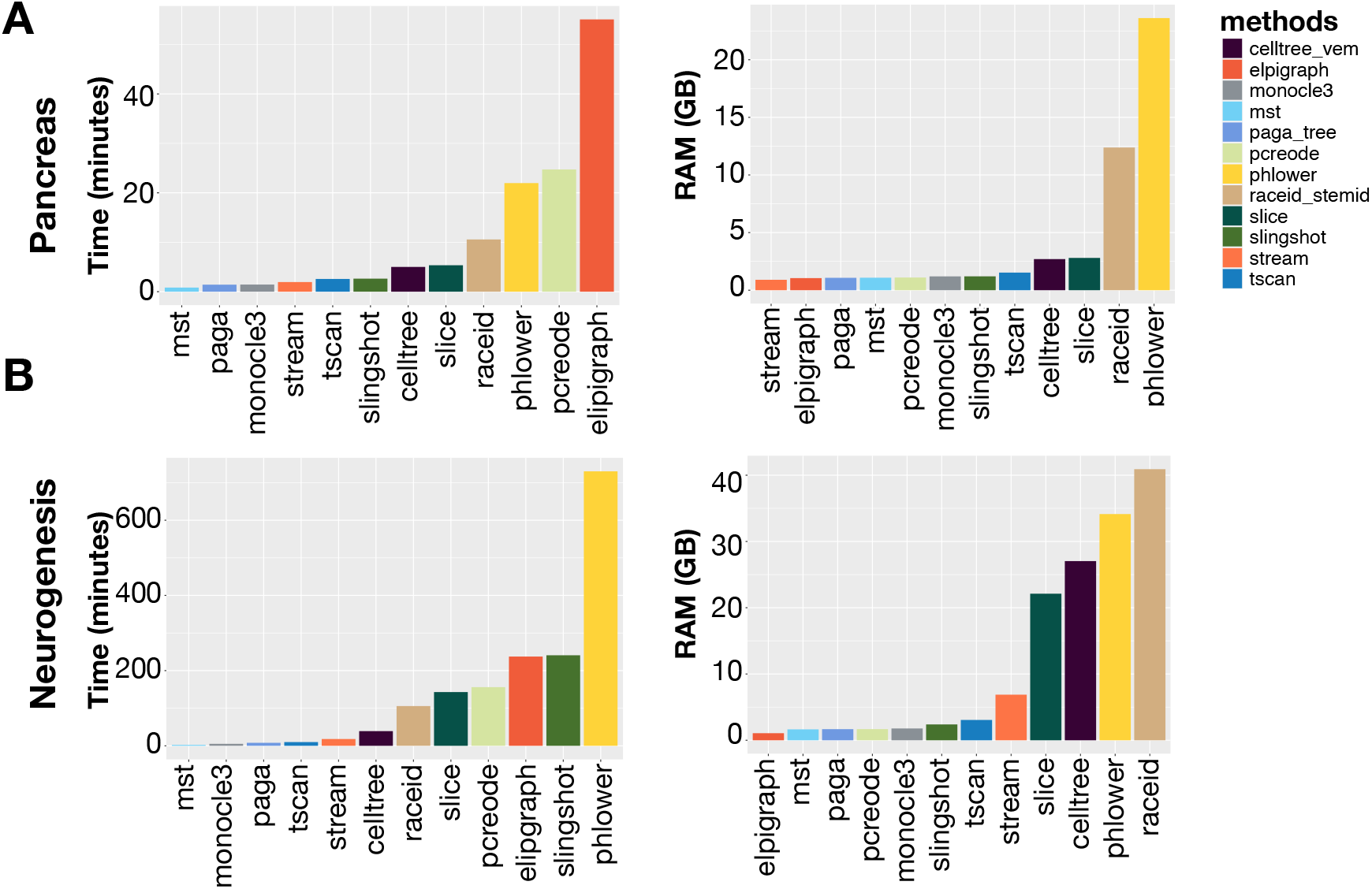
Time and Memory Benchmarking: **A)** We show the time (left) and memory (right) requirements for pancreas progenitor (3.7k cells) and **B)** neurogenesis (18k cells) data. The profiling was performed with the package memory-profiler (0.61.0) on a computer with an Intel i5 10400 processor (12 threads), 64GB RAM, running Linux Mint OS 21.1.

**Figure S5.**
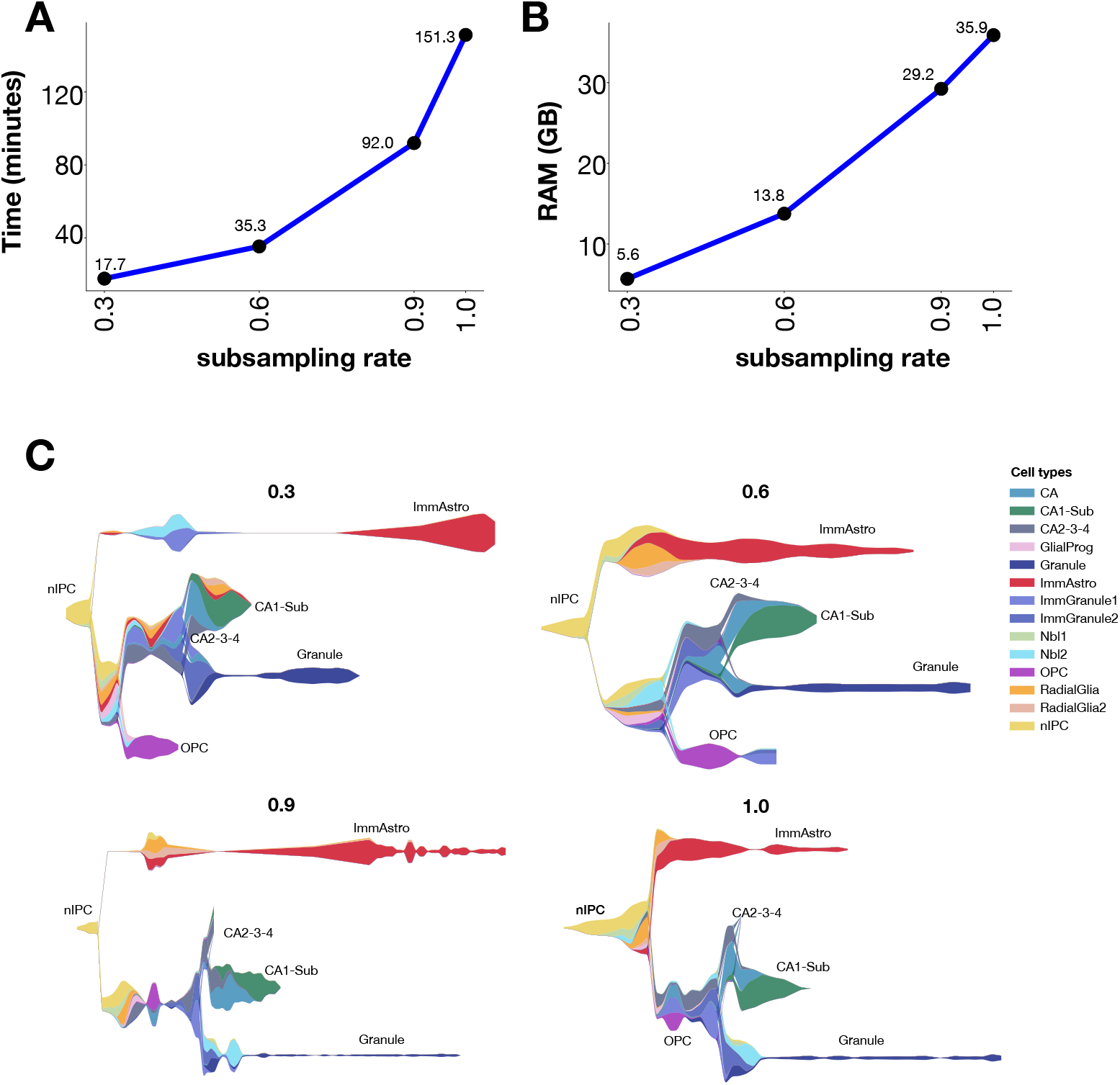
Neurogenesis data sub-sampling benchmark: **A-B)** We show the time and memory requirements of PHLOWER for the neurogenesis data (18k cells) after subsampling by using between 30% to 90% of cells. The use of half of the data provides a 8.6x speed up and requires 1/6 of the memory when compared to using all cells. Experiments were executed on a high perfomance computing node (AMD EPYC 7543 50/128 Cores 2.345G, 1024GB RAM, Rocky Linux 8.9.). Note that this provides an speed up of 4x compared to experiments in Supp. Fig. 5, which needed to be executed in a normal desktop computer due to super user requirements of Dynverse. **C)** PHLOWER stream trees obtained for the distinct sub-sampling procedures.

**Figure S6.**
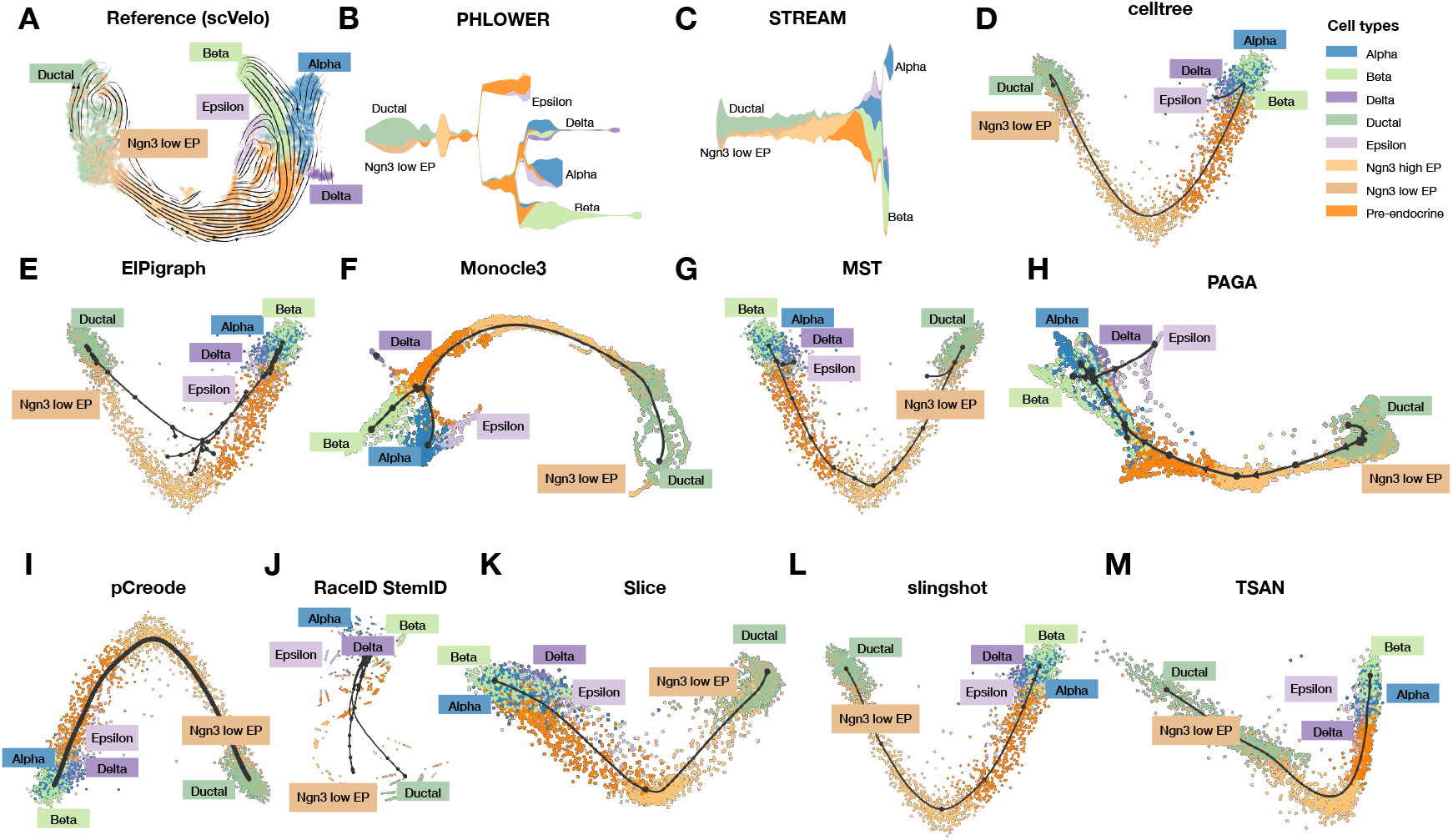
Pancreas trajectory for all evaluated methods. (A) Displays the original embedding of the pancreas progenitor data as reported in scVelo^38^. Colors corresponds to the cell annotation provided in^38^. This information is not provided to algorithms. B-M shows the results for all evaluated methods. We use the default embedding of the tool for visualization. For several approaches (Slingshot, MST, pCreode,CellTree and ElpiGraph), this is based on PCA as implemented in Dynverse.

**Figure S7.**
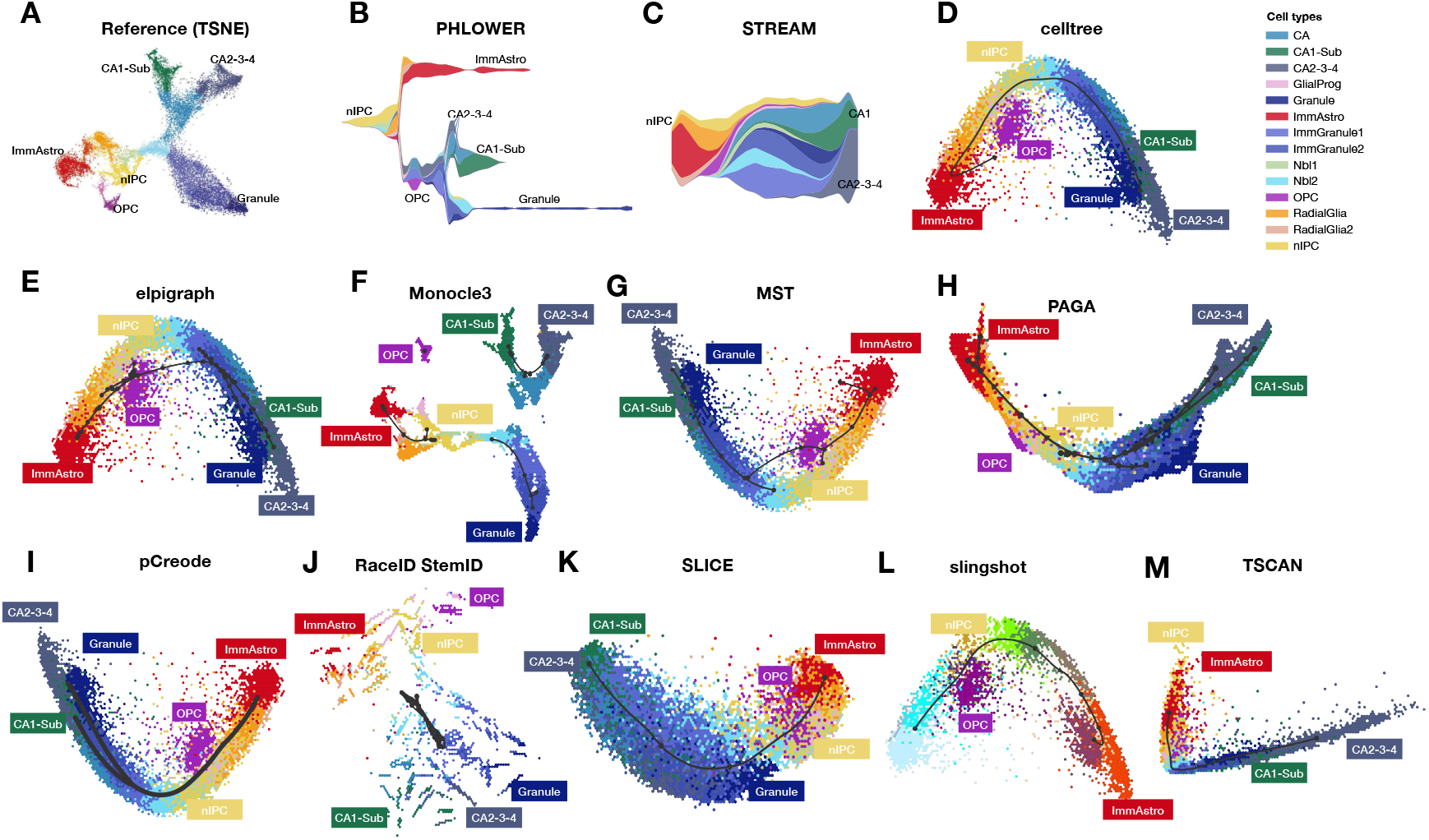
Neurogenesis trajectory for all evaluated methods. (A) Displays the original embedding of the neurogenesis data as reported in ^37^. Colors corresponds to the cell annotation provided in^37^. This information is not provided to algorithms. B-M shows the results for all evaluated methods.

**Figure S8.**
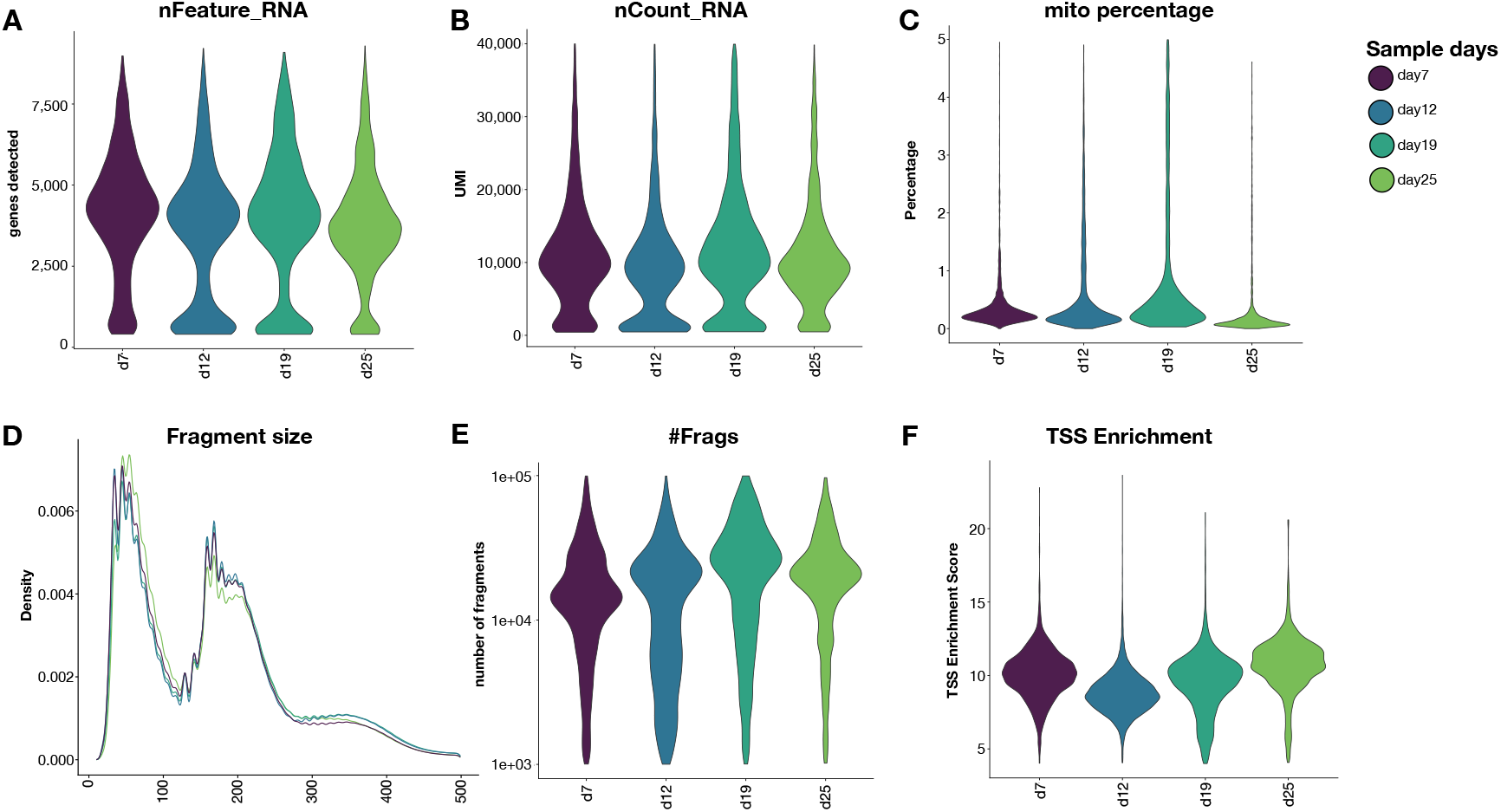
Quality check for kidney organoids single cell data. We show violin plots with quality check information after filtering and cell detection. These are: (A) number of features (genes) in RNA, (B) number of counts (transcripts) in RNA, (C) proportion of mitochondrial genes (RNA), (D) fragment sizes distribution (ATAC), number of fragments (ATAC) and (F) transcription start site enrichment. All libraries had similar values across days of sampling.

**Figure S9.**
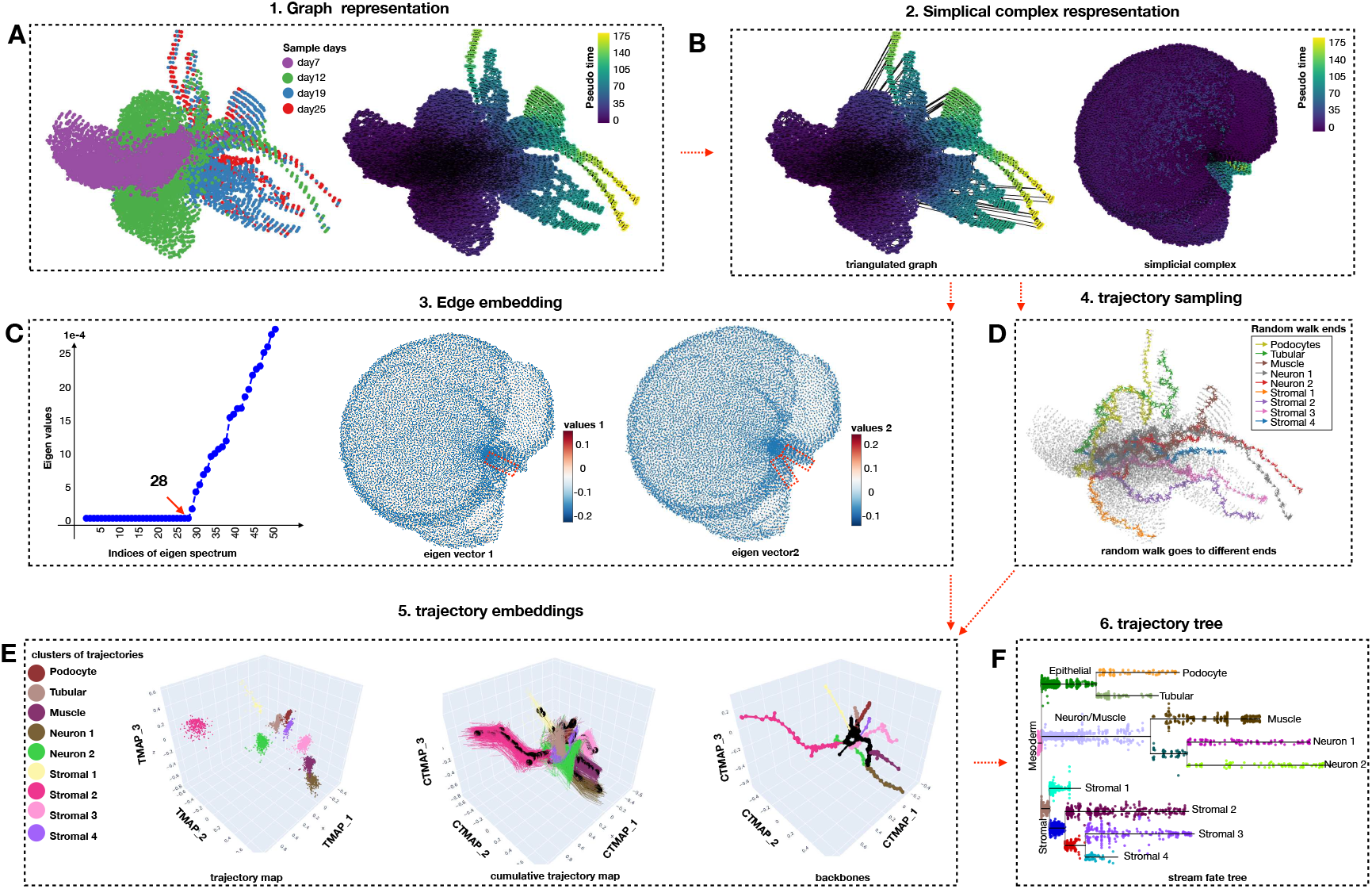
PHLOWER workflow for the kidney organoids data. Panels A-F are obtained as described in Supp. Fig. S1.

**Figure S10.**
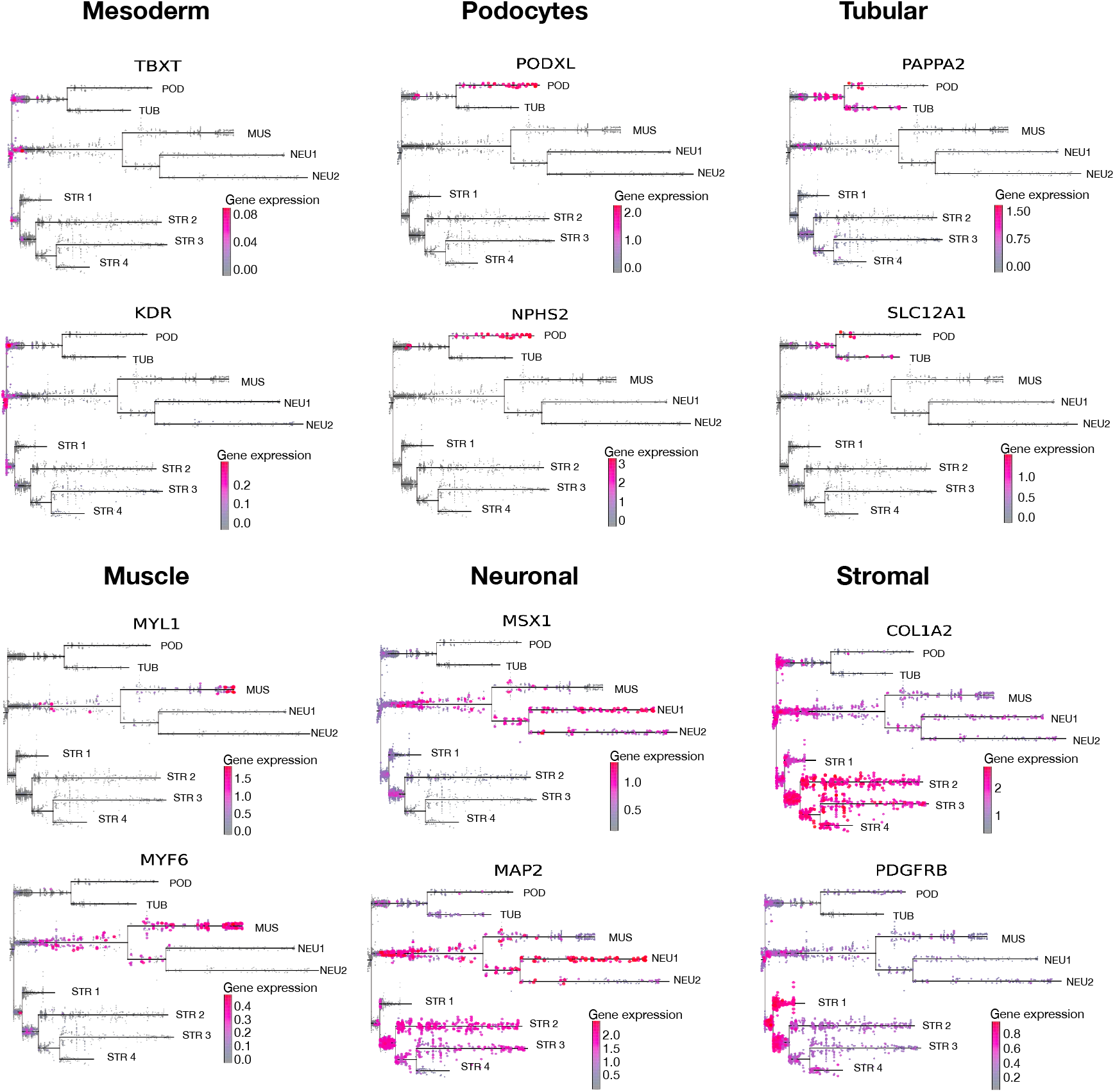
Gene markers for cell type of PHLOWER tree identification. **A)** Expressions of the gene markers TBXT, KDR, PODXL, NPHS2, PAPPA2, SLC12A1, MYL1, MYF6, MSX1, MAP2, COL1A2 and PDGFRB are shown in the PHLOWER tree (same genes as in Fig.3D).

**Figure S11.**
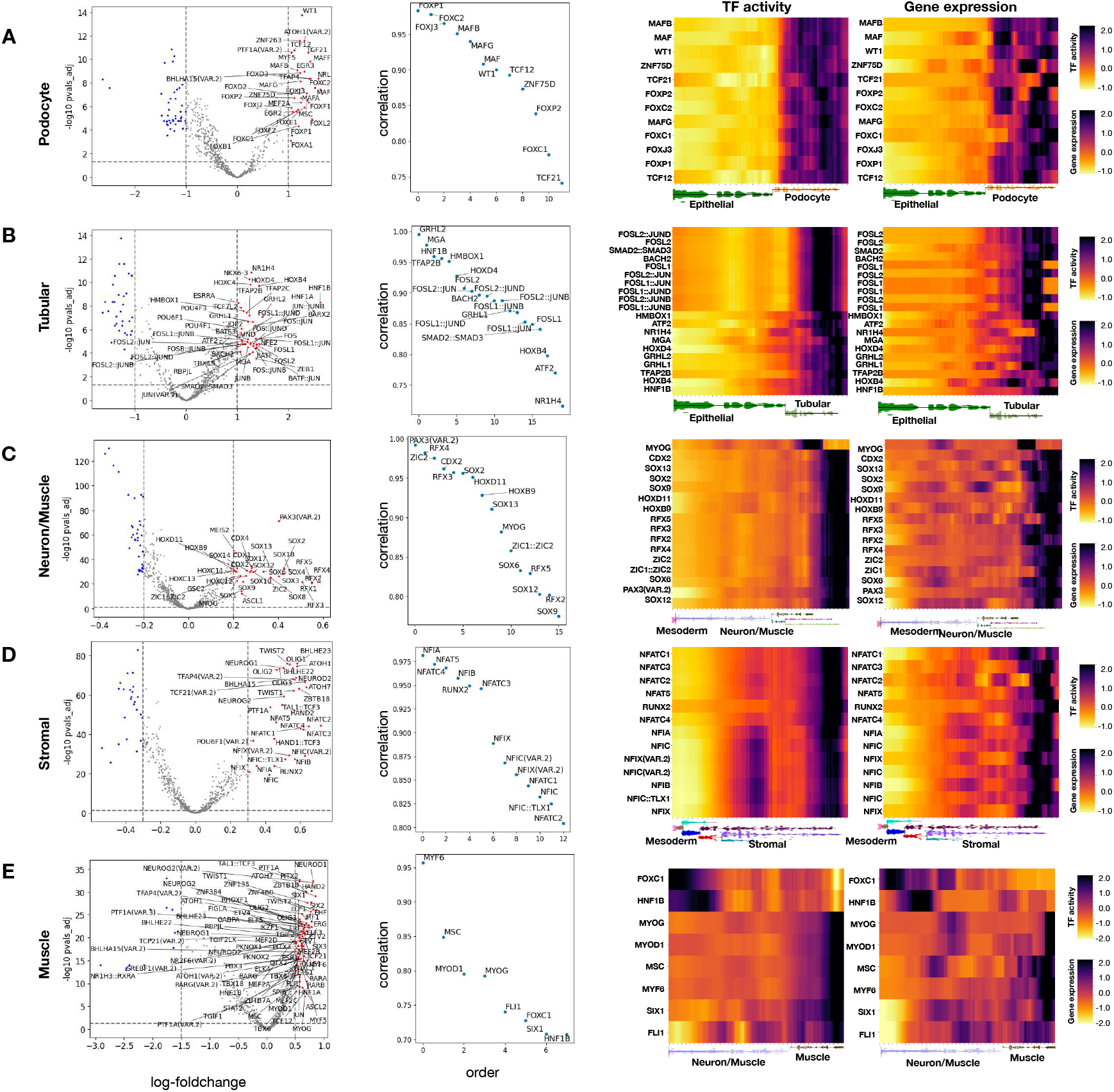
Regulators associated with cell branches. **A)** Regulators associated with Podocyte cells. We perform a differential expression analysis to find genes specific to podocytes (comparing with tubular cells) (left). Of these DE genes, we select transcription factors, which transcription factor activity (scATAC-seq), is highly correlated with gene expression (middle). Heatmaps display the TF activity and gene expression profiles of these TFs over the differentiation path from mesoderm towards podocyte cells. **B)** Same as (A) when contrasting tubular cells with podocytes. **C)** Sames as (A) when contrasting neuronal/muscle cells with all other branches. **D)** Same as (A) when contrasting stromal cells with all other branches. **E)** Same as (A) when contrasting Muscle cells with all other Neuron cells.

**Figure S12.**
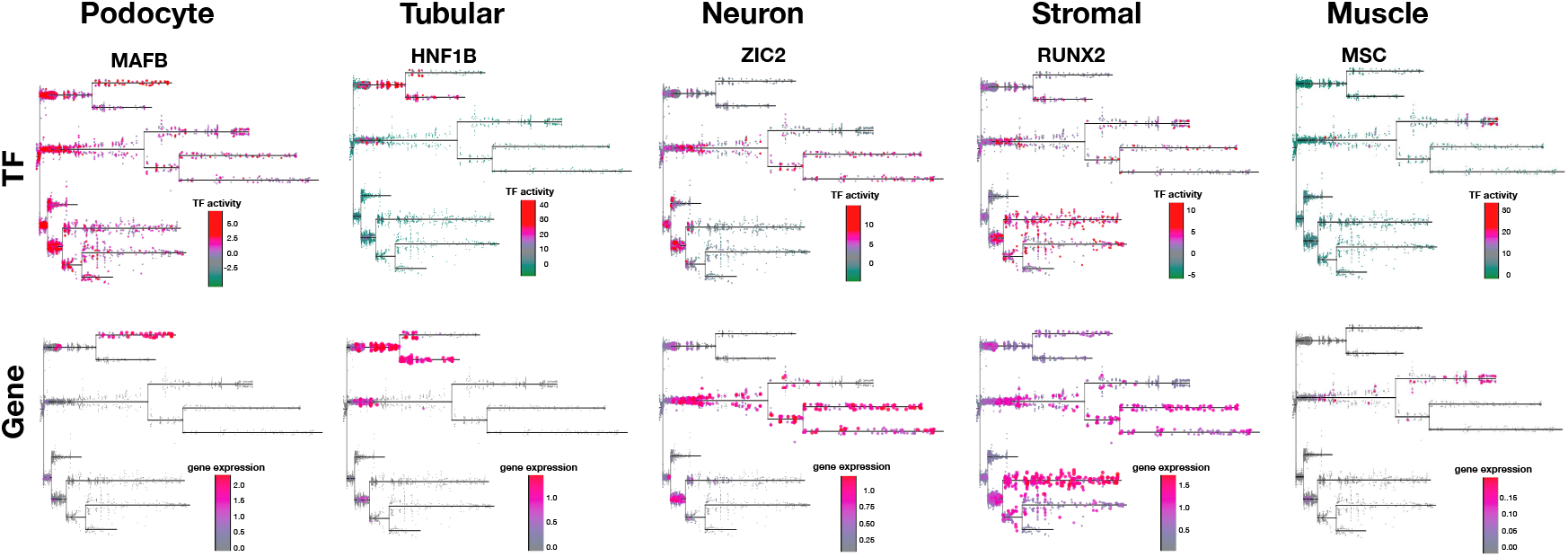
TF activity and gene expression showing in the PHLOWER estimated differentiation tree.

**Figure S13.**
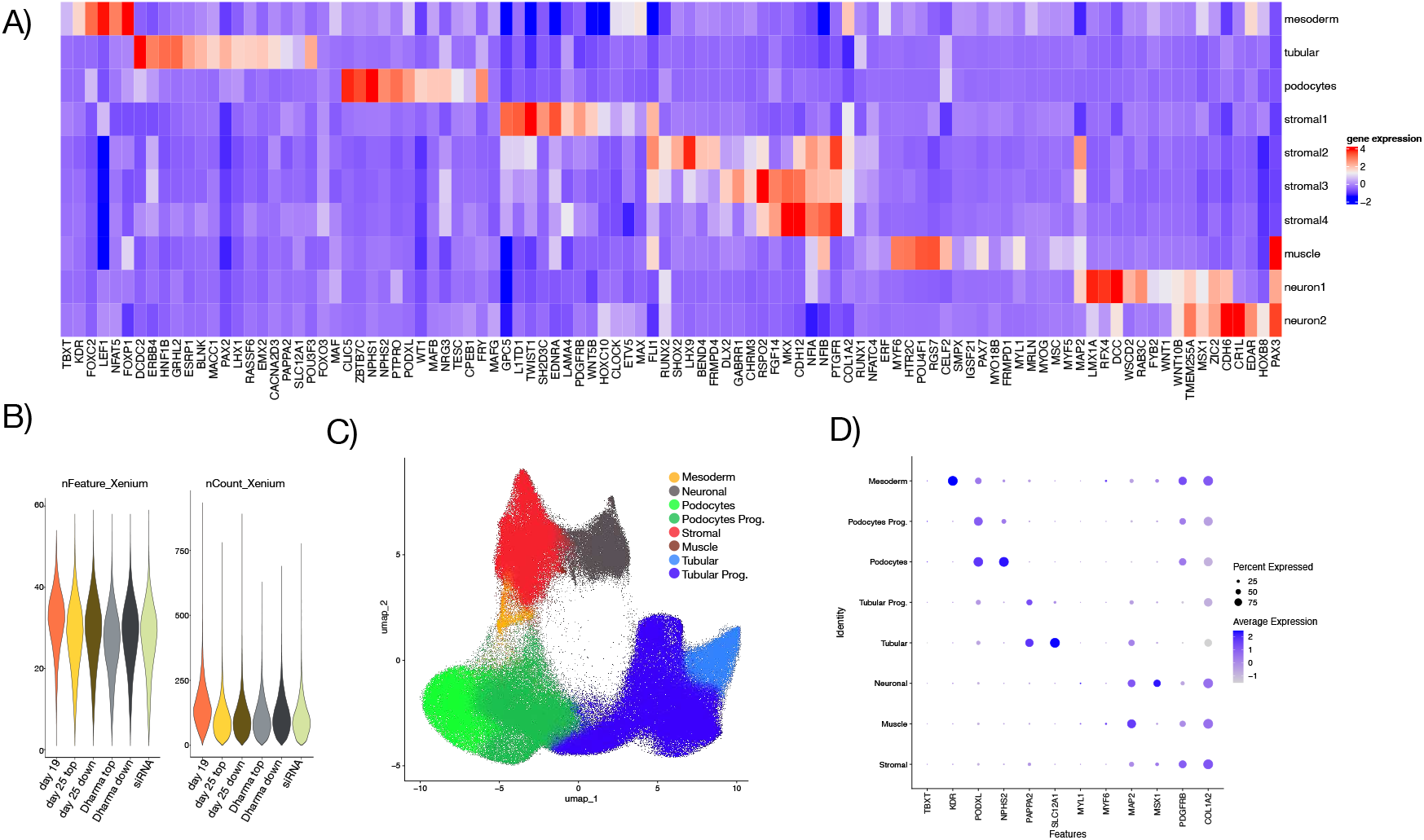
Xenium spatial profiling. **A)** Gene expression in multiome kidney organoid data, highlighting the 100 genes and transcription factors selected for the Xenium experiment. **B)** QC for the day 19, day 25, scrambled siRNA and siRNA treated spatial data. **C)** UMAP shows the cell types of the xenium data. **D)** Dot plots show the markers of cell types of the xenium data.

**Figure S14.**
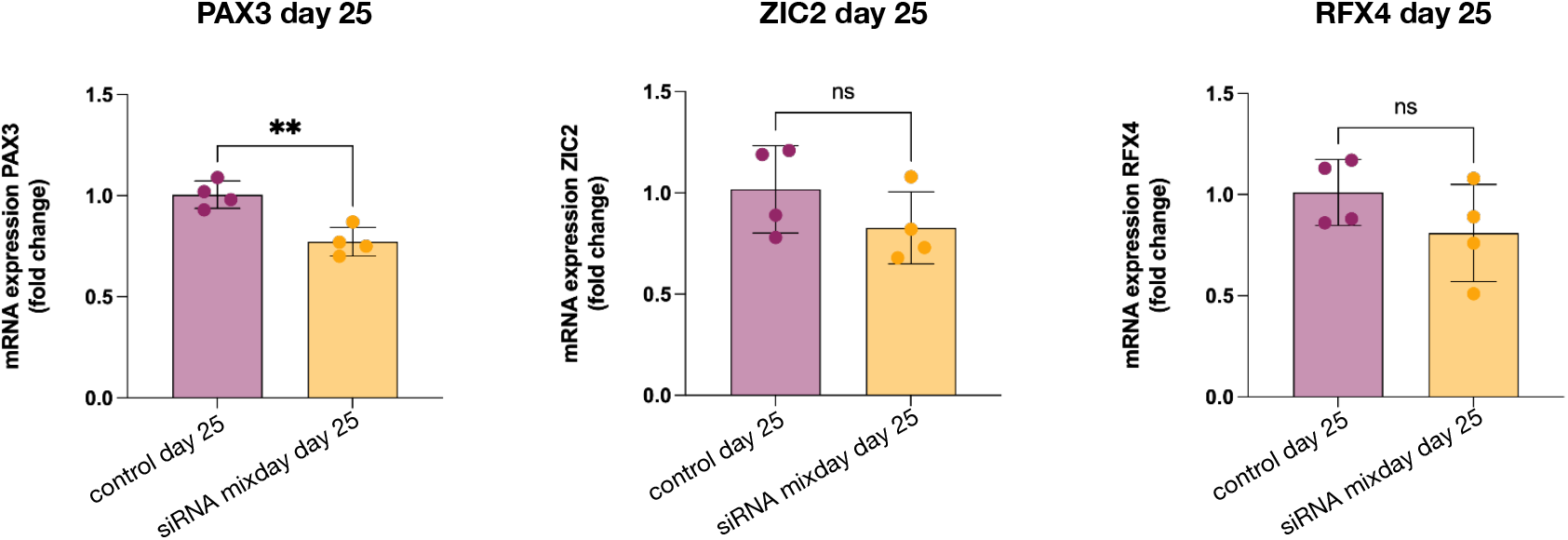
mRNA expression of genes targeted by siRNA experiments. Statistical test using unpaired *t*-test with bars representing mean ± sd; asterisks indicate P-values of <0.01 (**).

**Figure S15.**
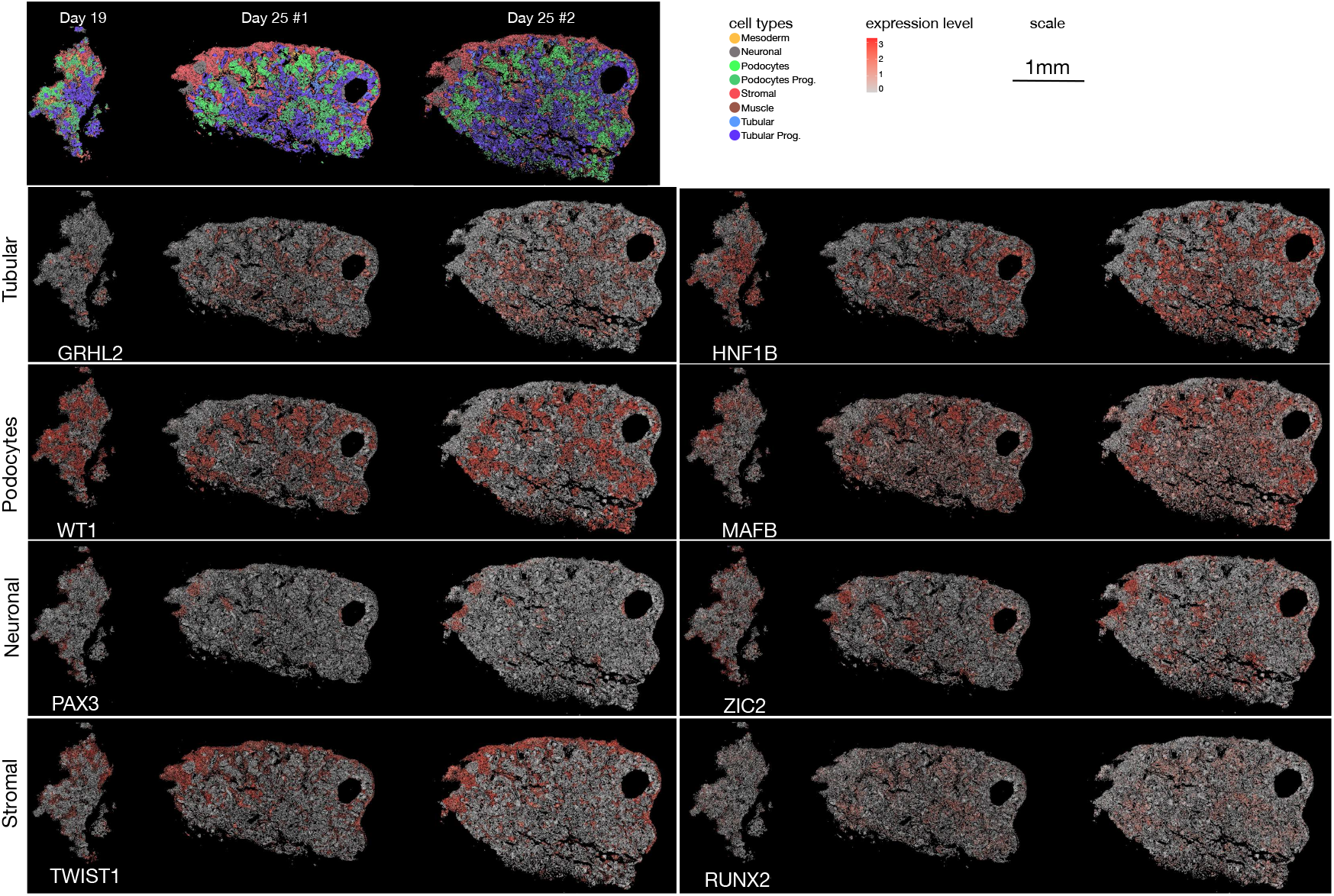
Spatial expression of cell type specific transcription factors for day 19 and day 25 kidney organoids.

**Figure S16.**
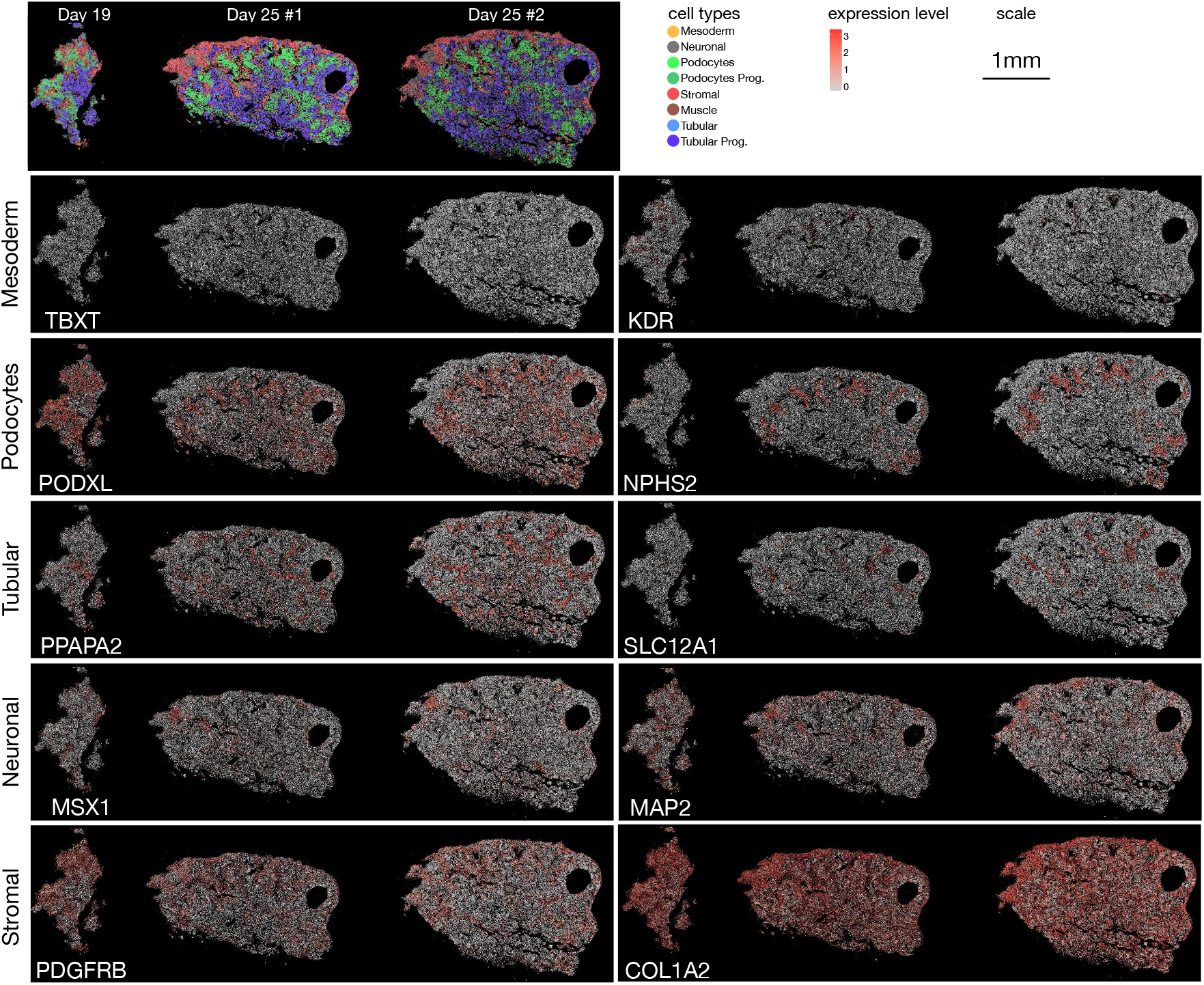
Spatial gene expression for cell type specific markers for day 19 and day 25 kidney organoids.

**Figure S17.**
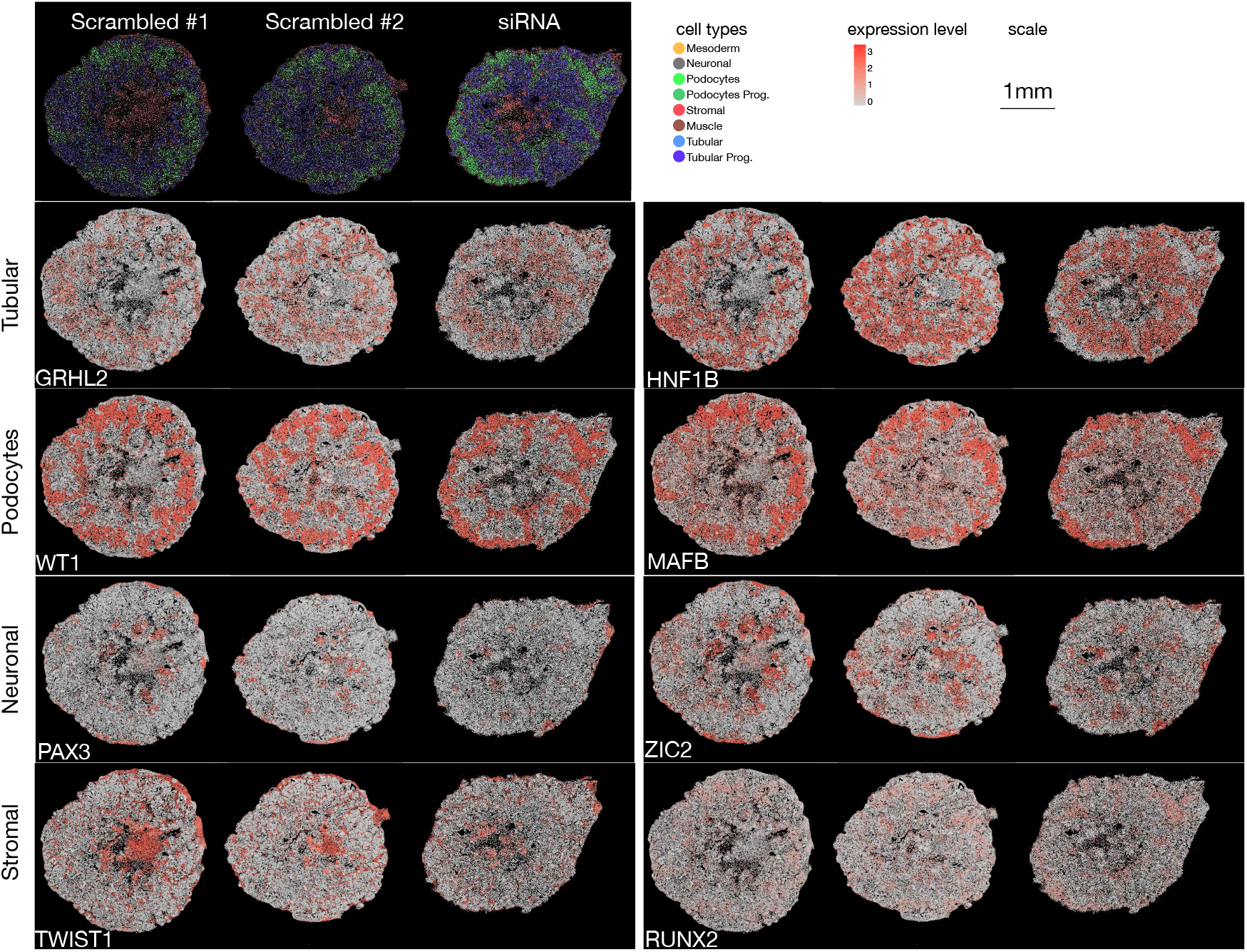
Spatial gene expression for cell type specific transcription factors for scrambled siRNA and siRNA treated.

**Figure S18.**
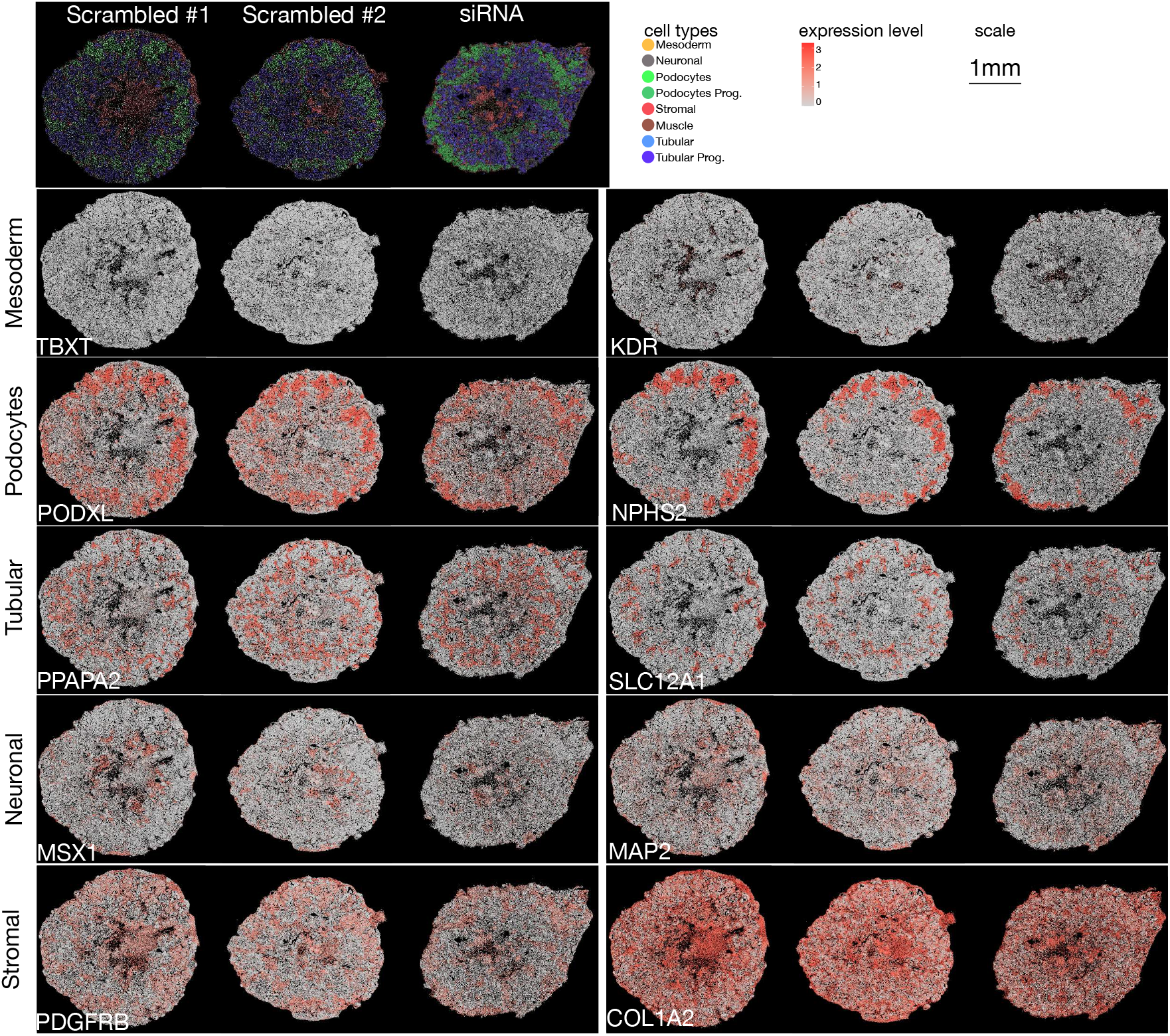
Spatial gene expression for cell type specific markers for scrambled siRNA and siRNA treated.

**Figure S19.**
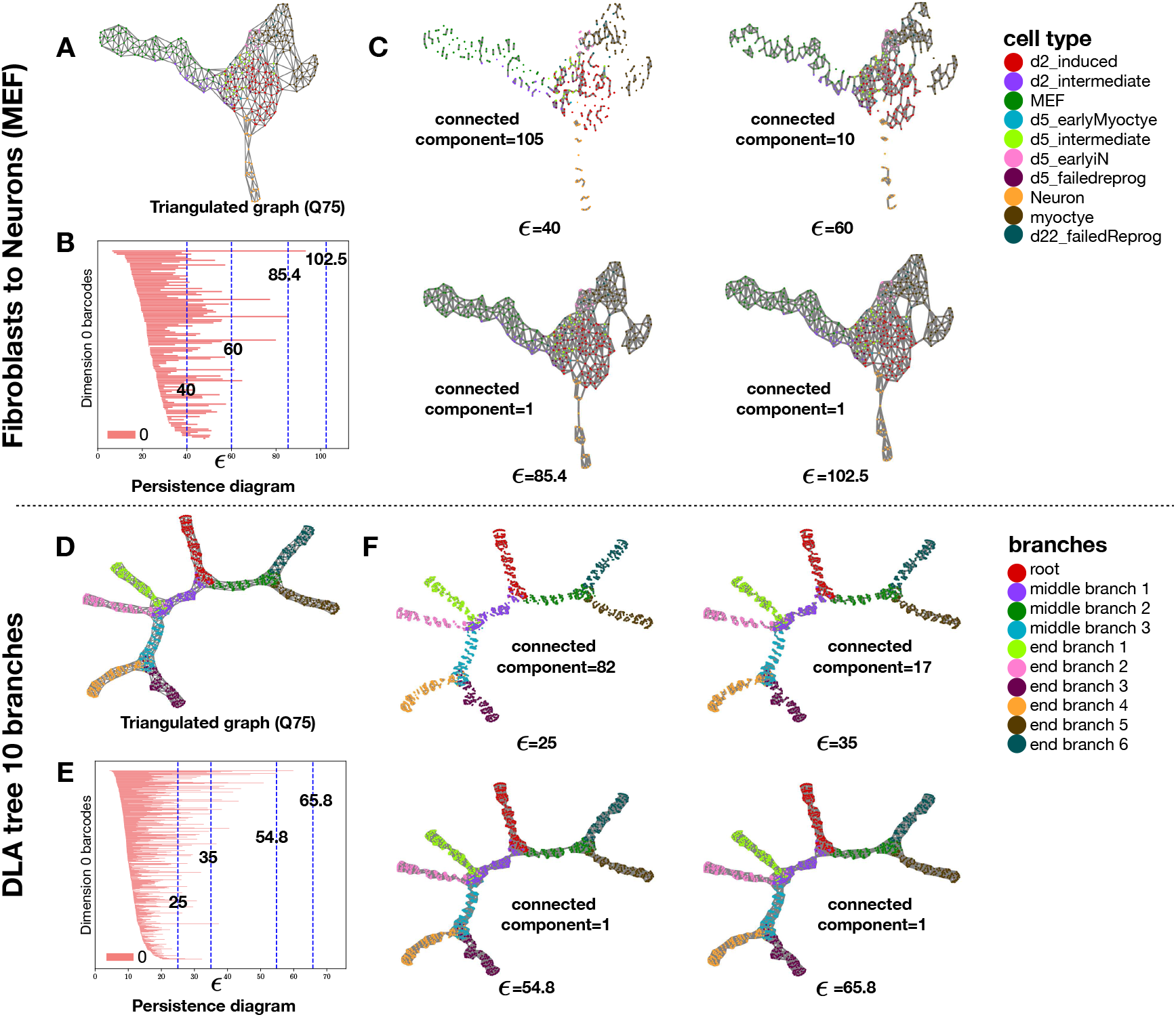
Persistent Homology Analysis. (A) We display the triangulated single cell graph (threshold *Q*_75_) for the MEF data. (B) Persistence diagram with 0-loops for the same data cloud from (A). (C) Graphs obtained after varying the the radius *ε* for several cutoffs values. We report the number of barcodes/connected components for each radius. The value 85.4 represents the first graph with a single connected component. Using an radius of factor 1.2 higher (102.5) leads to a graph with higher connectivity between nodes. This approach provides a graph similar to the filtering scheme used in PHLOWER (A). D-F same as A-C for the DLA simulated data with 10 branches. The python module gudhi (3.10.1) is used for the persistence analysis

**Figure S20.**
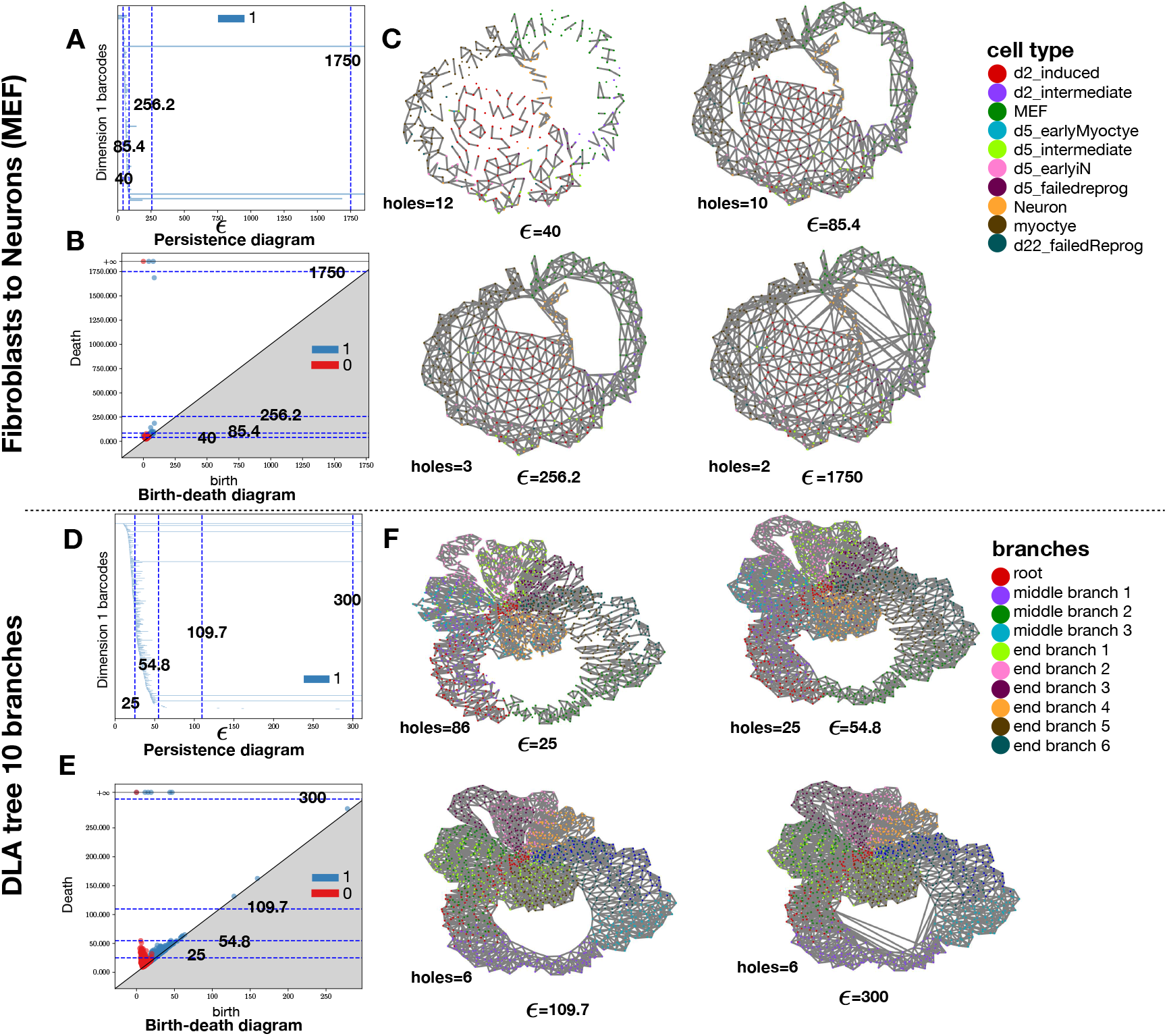
1-dimensional Persistent Homology. **(A)** Persistence diagram with 1-loops estimated on the SC of the MEF cells. (B) Birth and death diagram with both 0-loops (connected components) and 1-loops (holes) for the SC representing MEF cells. (C) SCs obtained after varying the the radius *ε* for several cutoffs values. Interestingly, the radius of 300, which is the lowest to remove all 1-loops but those with “infinity radius” find two holes as expected in this data set. D-F same as A-C for the simulated trees with 10 branches. In this more complex data set, the use of the threshold of 1750 provides a SC with 6 holes. This radius in the smallest radius such that only holes with “infinite size” are considered. These correspond to holes generated by artificially connecting cells with low and high pseudotime, precisely corresponding to features we want to extract with PHLOWER.

**Figure S21.**
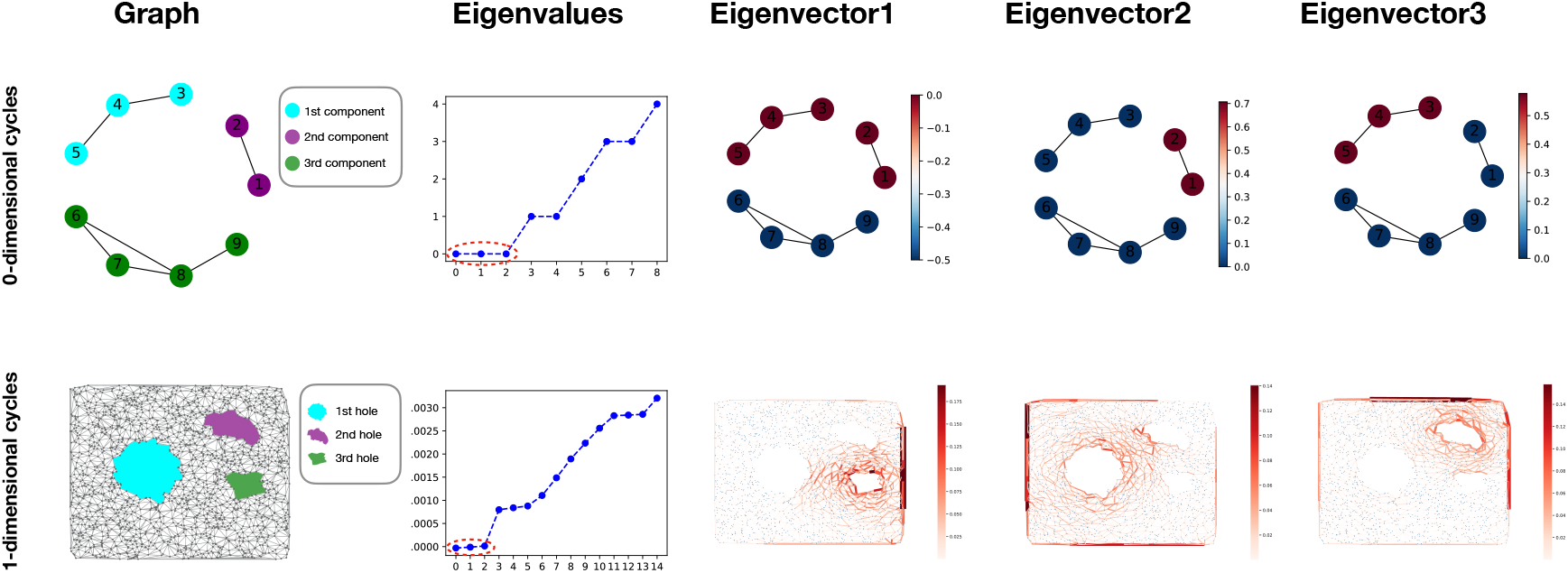
0-dimensional and 1-dimensional cycles. We demonstrate the detection of 0-dimensional cycles (connected components) and 1-dimensional cycles (holes) using the Graph Laplacian and the first-order Hodge Laplacian. In the first column, the upper side displays a graph with three 0-dimensional cycles (connected components), while the bottom side shows a simplicial complex with three 1-dimensional cycles (holes). In the second column, we observe the harmonic eigenvectors of the Hodge Laplacian on both the 1-simplicial complex (graph) and the 2-simplicial complex, each with three zero eigenvalues (up to numerical artifacts). The third, fourth and fifth columns depict these *harmonic* eigenvectors with zero-eigenvalues. We note that the eigenvectors of the graph Laplacian will be indexed over the nodes, whereas eigenvectors of the first Hodge Laplacian will be indexed over oriented edges. Signals of each eigenvector delineates nodes of distinct connected components (top) or contribution of edges to harmonic flows around holes (bottom) in the graph or simplicial complex.

**Figure S22.**
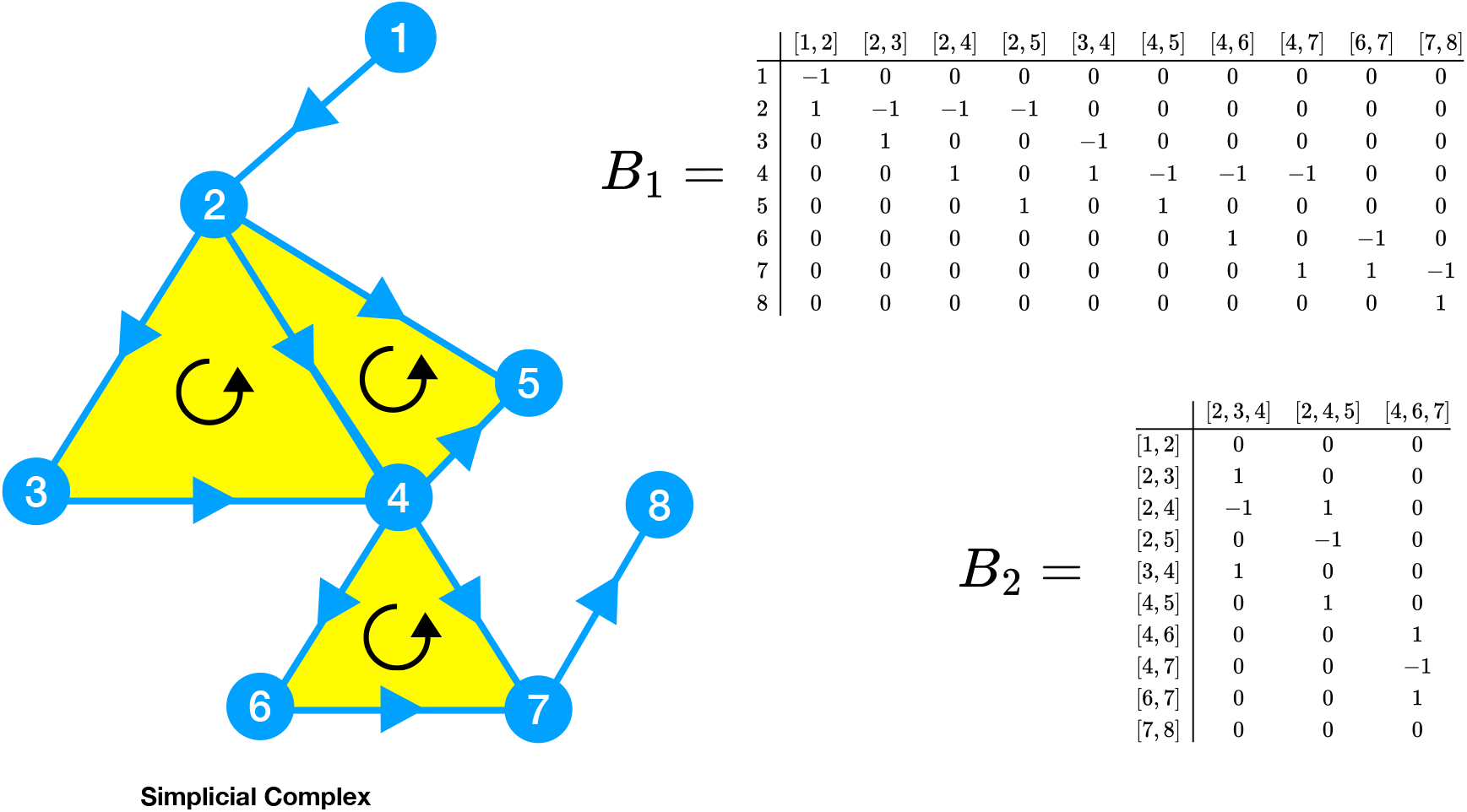
Example of an ordered simplicial complex and boundary maps. *B*_1_ **and** *B*_2_ **representing this complex**. Note that the order of edges/triangles correspond to a simple bookkeeping procedure, i.e. edge orientations are in the direction of increasing vertex index. The same applies for triangles.

**Figure S23.**
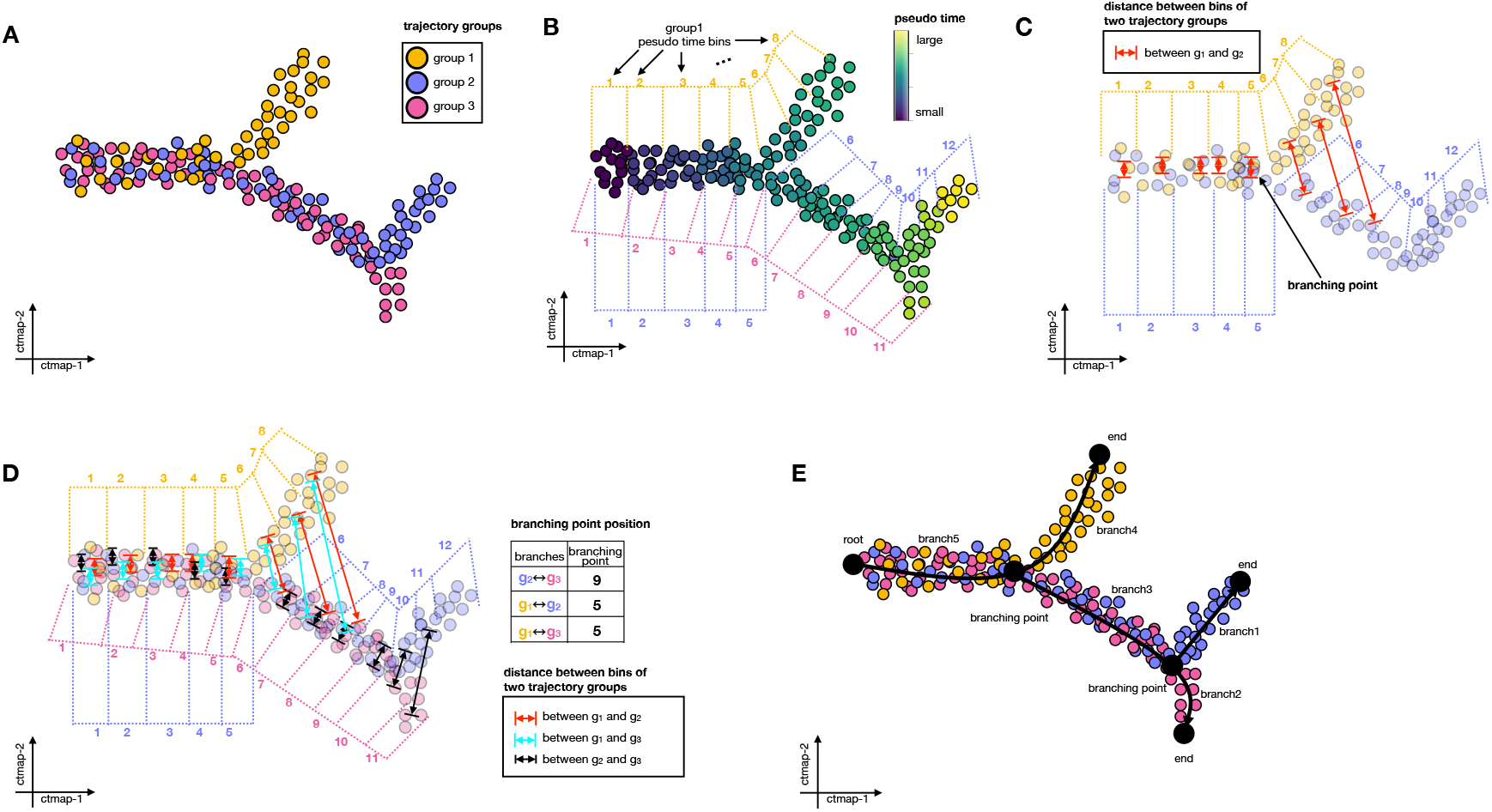
Schematic showing how to create a tree using edges lay in cumulative trajectory space. **A)** Example of three trajectory groups *g*_1_, *g*_2_, *g*_3_ each with a distinct color in the cumulative trajectory space. Every dot in this embedding represents an edge in the graph (or a cell differentiation event). **B)** PHLOWER takes the mean on the pseudo-time of the vertices (cells) connected by an edge and uses this as the edge pseudo-time estimate. PHLOWER next bins the edges by considering their pseudo-times. **C)** It then estimates average edge locations for every bin, which serves as a backbone for the trajectories. It also estimates the distance between edges in two groups but with the same bin index. **D)** Distance between bins are estimated for all groups and branchings points are detected whenever the distance between edges in two groups surpasses the distances of edges within a group for a given bin In the example, bin 9 is a branching point between groups 2 and 3 and bin 5 is a branching point between groups 1 and 2 and 1 and 3. **E)** To build the final tree, PHLOWER considers the branching points with the highest values towards the lowest values to build trees in a bottom-up approach, i.e. it merges trajectory 2 and 3 at bin 9. The next branching point is 5, which connects trajectory 1 with the tree formed by 2 and 3.

**Table S1.** List of all Dynverse data sets with a tree structure sorted by the number of branches. We only considered data sets (in bold) with at least 3 branches. Also, we excluded most “planaria” data sets as they represents sub-set of the **planaria-full_plass** data to avoid the introduction of a data set specific bias.

| name | nbranch | ncells | standard | nb/branch | criterion |
| --- | --- | --- | --- | --- | --- |
| **planaria-full_plass** | 26 | 18837 | silver | 724.5 | ✓ |
| planaria-pair-13_plass | 16 | 13265 | silver | 829.1 | ✗ repetitive |
| planaria-pair-9_plass | 13 | 11319 | silver | 870.7 | ✗ repetitive |
| planaria-pair-2_plass | 12 | 10653 | silver | 887.8 | ✗ repetitive |
| planaria-pair-12_plass | 11 | 11685 | silver | 1062.3 | ✗ repetitive |
| planaria-pair-7_plass | 10 | 11009 | silver | 1100.9 | ✗ repetitive |
| **ICM-monkey_nakamura** | 9 | 272 | silver | 30.2 | ✓ |
| planaria-pair-10_plass | 9 | 14636 | silver | 1626.2 | ✗ repetitive |
| planaria-pair-14_plass | 8 | 13281 | silver | 1660.1 | ✗ repetitive |
| planaria-pair-4_plass | 8 | 8960 | silver | 1120.0 | ✗ repetitive |
| **fetal-liver-fetal-hematopoiesis_mca** | 7 | 2559 | silver | 365.6 | ✓ |
| **oligodendrocyte-differentiation-subclusters_marques** | 7 | 4930 | silver | 704.3 | ✓ |
| planaria-pair-11_plass | 7 | 12978 | silver | 1854.0 | ✗ repetitive |
| **epiblast-monkey_nakamura** | 6 | 182 | silver | 30.3 | ✓ |
| **hematopoiesis-clusters_olsson** | 6 | 376 | silver | 62.7 | ✓ |
| **planaria-neuron-differentiation_plass** | 6 | 2349 | silver | 391.5 | ✓ |
| planaria-pair-3_plass | 6 | 8589 | silver | 1431.5 | ✗ repetitive |
| planaria-pair-6_plass | 6 | 8983 | silver | 1497.2 | ✗ repetitive |
| **embronic-mesenchyme-neuron-differentiation_mca** | 5 | 481 | silver | 96.2 | ✓ |
| **mesoderm-development_loh** | 4 | 504 | gold | 126.0 | ✓ |
| planaria-pair-5_plass | 4 | 12660 | silver | 3165.0 | ✗ repetitive |
| planaria-pair-8_plass | 4 | 9354 | silver | 2338.5 | ✗ repetitive |
| **planaria-parenchyme-differentiation_plass** | 4 | 1986 | silver | 496.5 | ✓ |
| **germline-human-both_guo** | 3 | 272 | gold | 90.7 | ✓ |
| **macrophage-salmonella_saliba** | 3 | 60 | gold | 20.0 | ✓ |
| **NKT-differentiation_engel** | 3 | 197 | gold | 65.7 | ✓ |
| **cortical-interneuron-differentiation_frazer** | 3 | 213 | silver | 71.0 | ✓ |
| **fibroblast-reprogramming_treutlein** | 3 | 355 | silver | 118.3 | ✓ |
| **hepatoblast-differentiation_yang** | 3 | 504 | silver | 168.0 | ✓ |
| **kidney-collecting-duct-subclusters_park** | 3 | 1533 | silver | 511.0 | ✓ |
| **neonatal-rib-cartilage_mca** | 3 | 2221 | silver | 740.3 | ✓ |
| **placenta-trophoblast-differentiation_mca** | 3 | 1001 | silver | 333.7 | ✓ |
| **trophoblast-stem-cell-trophoblast-differentiation_mca** | 3 | 19467 | silver | 6489.0 | ✓ |
| distal-lung-epithelium_treutlein | 2 | 59 | silver | 29.5 | ✗ simple |
| planaria-muscle-differentiation_plass | 2 | 2338 | silver | 1169.0 | ✗ simple |
| planaria-pair-1_plass | 2 | 10578 | silver | 5289.0 | ✗ simple |
| thymus-t-cell-differentiation_mca | 2 | 1607 | silver | 803.5 | ✗ simple |
| aging-hsc-old_kowalczyk | 1 | 873 | gold | 873.0 | ✗ simple |
| aging-hsc-young_kowalczyk | 1 | 493 | gold | 493.0 | ✗ simple |
| developing-dendritic-cells_schlitzer | 1 | 238 | gold | 238.0 | ✗ simple |
| germline-human-female_guo | 1 | 100 | gold | 100.0 | ✗ simple |
| germline-human-female-weeks_li | 1 | 666 | gold | 666.0 | ✗ simple |
| germline-human-male_guo | 1 | 166 | gold | 166.0 | ✗ simple |
| germline-human-male-weeks_li | 1 | 649 | gold | 649.0 | ✗ simple |
| hematopoiesis-gates_olsson | 1 | 316 | gold | 316.0 | ✗ simple |
| human-embryos_petropoulos | 1 | 1289 | gold | 1289.0 | ✗ simple |
| mESC-differentiation_hayashi | 1 | 414 | gold | 414.0 | ✗ simple |
| myoblast-differentiation_trapnell | 1 | 290 | gold | 290.0 | ✗ simple |
| pancreatic-alpha-cell-maturation_zhang | 1 | 322 | gold | 322.0 | ✗ simple |
| pancreatic-beta-cell-maturation_zhang | 1 | 562 | gold | 562.0 | ✗ simple |
| psc-astrocyte-maturation-glia_sloan | 1 | 455 | gold | 455.0 | ✗ simple |
| psc-astrocyte-maturation-neuron_sloan | 1 | 192 | gold | 192.0 | ✗ simple |
| stimulated-dendritic-cells-LPS_shalek | 1 | 540 | gold | 540.0 | ✗ simple |
| stimulated-dendritic-cells-PAM_shalek | 1 | 407 | gold | 407.0 | ✗ simple |
| stimulated-dendritic-cells-PIC_shalek | 1 | 433 | gold | 433.0 | ✗ simple |

**Table S2.**
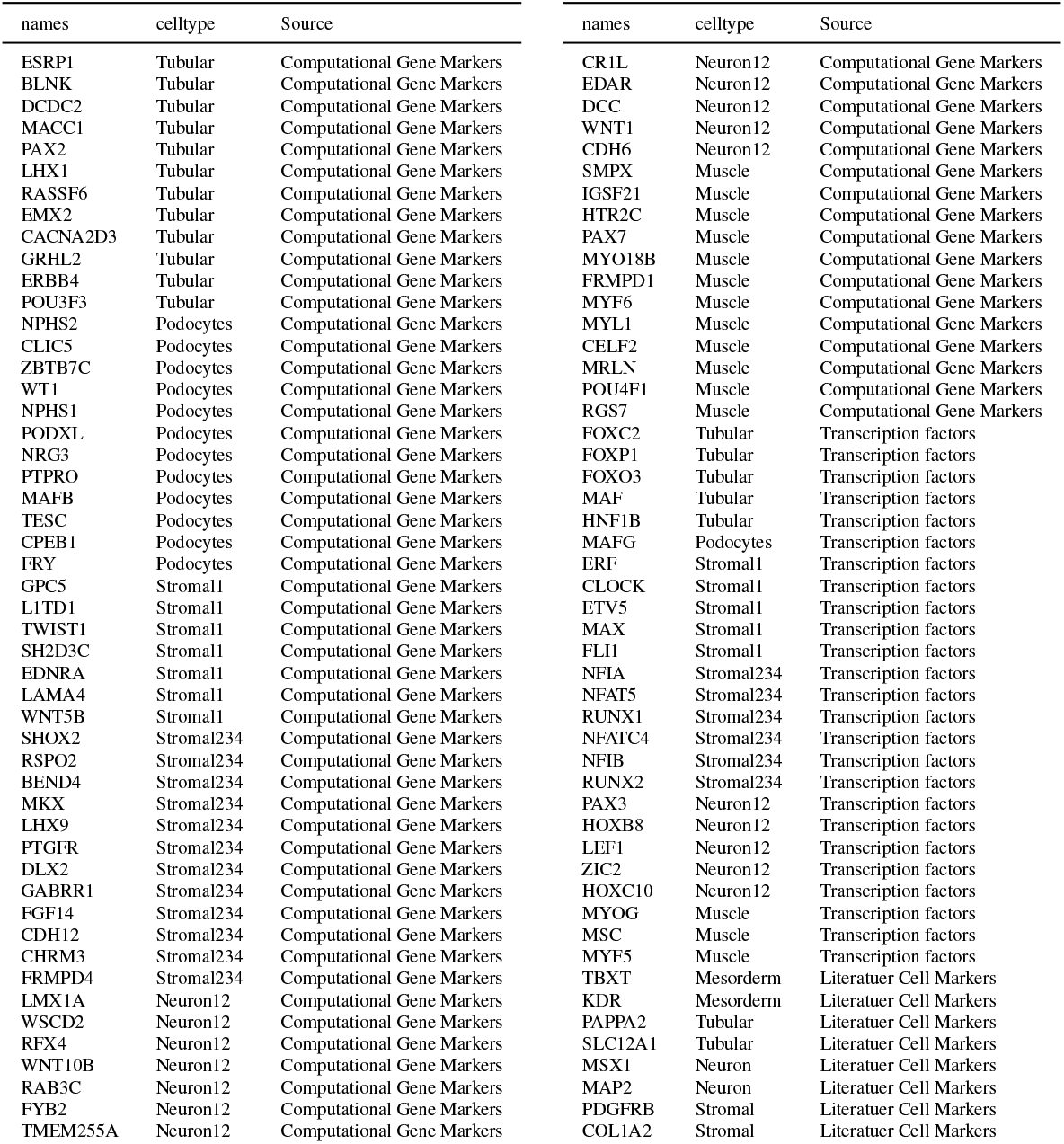
List of genes used in the Xenium spatial profiling panel. Source indicates if marker were based on computational markers from the mulitome data; literature markers or TFs predicted by PHLOWER and scMEGA.

## Footnotes

1 Note that some tools (Palantir and MIRA^7, 35^) do not infer differentiation trees and can therefore not be evaluated here.

